# A bHLH-PAS protein regulates light-dependent diurnal rhythmic processes in the marine diatom *Phaeodactylum tricornutum*

**DOI:** 10.1101/271445

**Authors:** Rossella Annunziata, Andrés Ritter, Antonio Emidio Fortunato, Soizic Cheminant-Navarro, Nicolas Agier, Marie J. J. Huysman, Per Winge, Atle Bones, François-Yves Bouget, Marco Cosentino Lagomarsino, Jean Pierre Bouly, Angela Falciatore

## Abstract

Periodic light-dark cycles govern the timing of basic biological processes in organisms inhabiting land as well as the sea, where life evolved. Although prominent marine phytoplanktonic organisms such as diatoms show robust diurnal rhythms in growth, cell cycle and gene expression, the molecular foundations controlling these processes are still obscure. By exploring the regulatory landscape of diatom diurnal rhythms, we unveil the function of a *Phaeodactylum tricornutum* bHLH-PAS protein, *Pt*bHLH1a, in the regulation of light-dependent diurnal rhythms. Peak expression of *PtbHLH1a* mRNA occurs toward the end of the light period and it adjusts to photoperiod changes. Ectopic over-expression of *Pt*bHLH1a results in lines showing a phase shift in diurnal cell fluorescence, compared to the wild-type cells, and with altered cell cycle progression and gene expression. Reduced oscillations in gene expression are also observed in overexpression lines compared to wild-type in continuous darkness, showing that the regulation of rhythmicity by *Pt*bHLH1a is not directly dependent on light inputs and cell division. *Pt*bHLH1a homologs are widespread in diatom genomes which may indicate a common function in many species. This study adds new elements to understand diatom biology and ecology and offers new perspectives to elucidate timekeeping mechanisms in marine organisms belonging to a major, but underinvestigated branch of the tree of life.

**SIGNIFICANCE STATEMENT:** Most organisms experience diurnal light-dark changes and show rhythms of basic biological processes such that they occur at optimal times of the day. The ocean harbours a huge diversity of organisms showing light-dependent rhythms, but their molecular foundations are still largely unknown. In this study, we discover a novel protein, *Pt*bHLH1a that regulates cell division, gene expression and the diurnal timing of these events in the marine diatom *Phaedoactylum tricornutum*. The identification of *PtbHLH1a*-like genes in many diatom species suggests a conserved function in diurnal rhythm regulation in the most species-rich group of algae in the ocean. This study unveils critical features of diatom biology and advances the field of marine rhythms and their environmental regulation.

## INTRODUCTION

The Earth’s rotation means that life evolved under a 24h diurnal cycle of alternate light and dark periods. Most living organisms have developed daily rhythms of many fundamental biological processes, ranging from physiology to behaviour, such that they occur at optimal times of the day (1) which can enhance fitness (2, 3). These rhythms are the product of the coordinated action of signals from endogenous timekeepers, together with environmental and metabolic inputs (4, 5). Robust diel rhythms in growth, cell cycle, gene expression, pigment synthesis, phototactic movements and bioluminescence have been also observed in a variety of phytoplanktonic organisms, including diatoms (6-13). Diatoms represent the most species-rich group of algae in the ocean and populate a wide range of aquatic environments (14, 15). These algae of the Stramenopile lineage show peculiar genomic, metabolic and cellular features, and are evolutionarily distant from the most studied model organisms in the field of biological rhythms (16-20). Diatoms have an impressive capacity to deal with environmental changes thanks to sophisticated acclimation mechanisms (21-25). Recent genome-wide analyses also showed that 25% of the diurnal transcriptome is influenced by light-dark cycles in the centric diatom *Thalasiossira pseudonana* (26). Moreover, detailed diurnal studies in the pennate diatom *Phaeodactylum tricornutum* highlighted a strict temporal separation of transcriptional gene networks (27, 28), as previously observed in other algae (29). Tight diurnal control of the *P*. *tricornutum* cell cycle has also been observed (27, 30, 31), with light onset triggering cell cycle progression through the transcriptional activation of the diatom-specific cyclin dsCYC2 by the blue light sensor Aureochrome-1a (30). Furthermore, the transcription factor bZIP14 has recently been identified as a diurnal activator of the tricarboxylic acid (TCA) cycle, a process restricted to the late afternoon in diatoms (32). Together, these studies illustrate complex regulation of diurnal cellular activities in *P. tricornutum*. However, the molecular mechanisms orchestrating diurnal processes are still unknown in diatoms and many other phytoplanktonic organisms. Notably, no orthologs of the circadian clock components discovered in bacteria, fungi, animals or plants have been found in the diatom genomes except for cryptochromes (33). Nonetheless, a number of proteins containing bHLH-PAS domains, which feature in genes involved in the regulation of rhythmic processes in animals (34), have been identified in diatom genomes (35).

In this work, we integrate transcriptomic, physiological and functional analyses to explore the regulatory landscape of *P. tricornutum* diurnal rhythms. We uncover the function of a bHLH-PAS protein (*Pt*bHLH1a) in the regulation of critical light-dependent rhythmic processes, such as cell cycle and diurnal transcription. Phylogenetic analyses reveal that bHLH1a homologs are widely distributed in diatoms, thus we speculate a common function in many diatom species. These results open the way to new exploration of diatom genomes in search of their elusive molecular timekeepers.

## RESULTS

### Transcriptome profiling identifies potential regulators of diurnal rhythms in *P. tricornutum*

To identify potential regulators of cellular rhythmicity in *P. tricornutum*, a publicly available diurnal transcriptomic dataset (27) was analyzed. One hundred and four genes with robust diel oscillating expression were selected, of which eight were photoreceptors (30, 33, 36, 37) and 66 were transcription factors (TFs) (35), which might be involved in the generation of light-dependent rhythms. The remaining 30 genes selected were potential output genes implicated in diel rhythmic processes (pigment synthesis, cell cycle regulation and photosynthesis) (Table S1). The transcriptional dynamics of the selected genes were examined in a 16:8-h light:dark (L:D) photocycle for 32h. Hierarchical clustering (HCL) analysis of the resulting expression profiles revealed 4 clusters of co-expressed genes, termed A-D (Fig. 1A), with peak expression at different times between dawn and dusk (Fig. 1B). Cluster A phased at dawn, suggesting a transcriptional anticipation of the light onset (Fig. 1A-B). This cluster comprised 18 genes including 14 TFs, mostly belonging to the Heat Shock Transcription Factor family (HSF), two DNA repair enzymes CPD photolyases (*CPD2* and *CPD3*) and one carotenoid synthesis enzyme (*PDS1*). Cluster B phased around 7h Zeitgeber Time (hours after illumination, ZT) and encompassed 36 genes (Fig. 1A-B), including the ds*CYC2* gene controlling the onset of cell division (30). Cluster B also contained 18 TFs, of which eight were sigma factors putatively involved in the regulation of chloroplast transcription, three genes implicated in photoprotection (*LHCX1*, *Vdl2* and *Zep1*) and the chlorophyll synthesis *POR1* gene. Such active transcription of genes involved in chloroplast activity during the light period has been shown previously (21, 38). The blue light sensors *Aurochrome1b* and the cryptochromes *CPF1* and *CryP-like* also belonged to cluster B and show a strong expression following light onset, in accordance with previous observations (39-41). Cluster C phased around ZT9 (Fig. 1A-B) and comprises 9 TFs and 10 metabolism-related genes, including genes encoding photosynthetic apparatus. Finally, cluster D phased before dusk and included 23 TF genes including the TCA cycle regulator *bZIP14* (27, 32), likely contributing to preparing cells for light to dark transition (Fig. 1A-B). Cluster D also contained the *CPF4* and the Far-Red light sensing phytochrome (*DPH1*) whose peak expression at the end of the light period has been observed previously (36, 39).

**Fig. 1.**
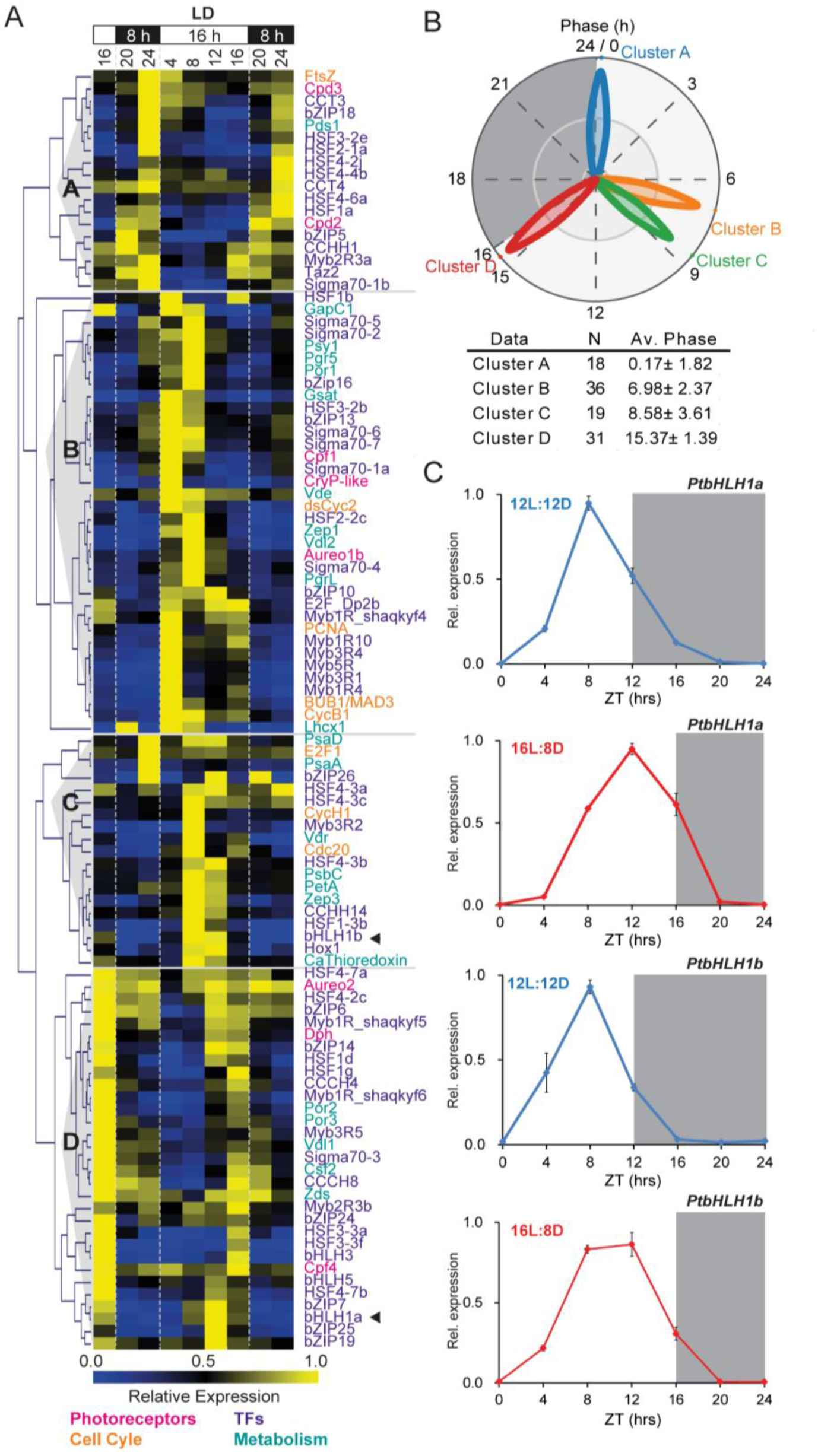
Diurnal expression analysis of selected rhythmic *P. tricornutum* genes. A) nCounter expression analysis of 104 selected genes in cells grown under a 16L:8D photoperiod and sampled every 4 hours for 32 hours. Four major groups of co-regulated genes (A-D) are shown based on hierarchical clustering. Expression values were normalized using *PtRPS*, *PtTBP* and *PtACTIN12* as reference genes and represent the average of three biological replicates. The *PtbHLH1a* and *PtbHLH1b* gene expression profiles are indicated with arrowheads. B) Polar plot and table showing the average phases of expression (intended as expression peaks during the analyzed period) of the four gene clusters, calculated using the MFourfit averaged method. Plot petal length is proportional to the standard deviation of phases over biological replicates. In the table, N: number of genes within each cluster; Av. Phase: average periods and phases of expression in hours. C) Diurnal expression profiles of *PtbHLH1a* and *PtbHLH1b* in Wt cells grown under 12L:12D (blue line) and 16L:8D (red line) photocycles analysed by qRT-PCR. Expression values were normalized using *PtRPS* and *PtTBP* as reference genes and represent the average of three biological replicates ± s.e.m (standard error of the mean; black bars). For each gene, the expression value is relative to its maximum expression (maximum expression=1). Light and dark periods are represented by white and grey regions respectively.

Together these results underline the existence of tight transcriptional programs phasing at discrete moments of the day which potentially control the timing of cellular activities along the diurnal cycle.

### *PtbHLH1a* expression is adjusted in a photoperiod-dependent manner

Our analysis identified two TFs, *Pt*bHLH1a (Phatr3_J44962) belonging to cluster D and *Pt*bHLH1b (Phatr3_J44963) belonging to cluster C, which each have a Per-ARNT-Sim (PAS) domain in conjunction with a bHLH DNA-binding domain. Because bHLH-PAS proteins have been shown to be involved in the regulation of rhythmic processes in animals (4, 34, 42), the expression profiles of *PtbHLH1a* and *PtbHLH1b* were examined in *P. tricornutum* cells growing under different photoperiods. *PtbHLH1a* expression peaked at ZT8 in the 12L:12D photoperiod and at ZT12 in the 16L:8D photoperiod, 4 hours before the end of the light period in both cases, then gradually decreased to below detection limits at ZT0 (Fig. 1C). Transcription of *PtbHLH1b* appeared to start earlier than that of *PtbHLH1a*. In cells entrained in 12L:12D cycles, *PtbHLH1b* expression peaked at ZT8, whereas it peaked between ZT8/ZT12 in 16L:8D photoperiods (Fig. 1C). Thus, *PtbHLH1b* expression onset almost coincided in the two photoperiods although transcription dramatically dropped after ZT8 in 12L:12D, while remained at maximum levels up to ZT12 in long days (Fig. 1C).

The robustness of *PtbHLH1a* and *PtbHLH1b* diurnal expression profiles was further examined under stress conditions using recent transcriptome datasets from *P. tricornutum* cells grown in 12L:12D cycles in iron replete and deplete conditions (28). Iron homeostasis is diurnally regulated in phytoplankton (43) and it affects rhythmic processes such as cell cycle progression and diurnal gene expression in *P. tricornutum* (28). Interestingly, *PtbHLH1a* and *PtbHLH1b* expression profiles showed similar patterns in both control and iron depleted conditions, with peaks of expression before dusk at ZT9 (Fig. S1), similar to our observations (Fig. 1C).

Altogether these results demonstrate robust control of *PtbHLH1a* and *PtbHLH1b* diurnal expression timing, which is adjusted in a photoperiod-dependent manner and unaffected by iron depletion. The involvement of *Pt*bHLH1a in the regulation of diurnal light-dependent rhythmic processes was hypothesized considering a possible role in dusk anticipation.

### *PtbHLH1a* ectopic expression determines phase shifts in cellular rhythmicity

To determine *Pt*bHLH1a’s function, cell lines were generated expressing HA-tagged bHLH1a under the regulation of the *Light harvesting complex protein family F2* promoter (*Lhcf2p*) (Fig. 2A), which activates transcription 3h after light onset (44). Gene expression analysis allowed the selection of three independent lines, hereafter named OE-1, OE-2 and OE-3, showing over-expression of the *PtbHLH1a* gene (Fig. 2B) and earlier expression timing compared to the wild type (Wt) strain (Fig. 4A and S2). Next, daily cellular rhythmicity was analyzed using the flow cytometer channel FL3 (excitation 488nm, emission 655-730 nm) that estimates chlorophyll a cellular content (8, 38), over the course of three days (Fig 2C). Cellular fluorescence displayed highly oscillating rhythms in 16L:8D grown cultures with a periodicity of approximately 24h (Fig. 2C, Table S2). Cell fluorescence in Wt cultures increased during daytime to peak around ZT13 (Fig. 2D and Table S2), and then started to decrease before night onset. This fall in fluorescence was concomitant with an increase in cell concentration ((38) and Fig. S4), likely reflecting chloroplast partitioning to daughter cells during cell division (7). Fluorescence progressively declined during the night period, reaching a trough in the early morning (Fig. 2C). Despite maintaining rhythmicity in the cellular fluorescence dynamics, OE-1 (Fig 2C), as well as OE-2 and OE-3 lines (Fig. 2D), displayed a remarkable phase shift of around 1-2 h in the maximum fluorescence timing compared to the Wt (Table S2). Cellular fluorescence phase responses were further investigated in resetting experiments. Wt and OE-1 cultures, that show the strongest phase shift phenotype, were grown in 16L:8D photoperiods, then transferred to 8L:16D and monitored for another 6 days. After the transfer to 8L:16D photocycles, the timing of maximum cell fluorescence in Wt cells was maintained for two days and then re-synchronized to the newly imposed photoperiod, peaking at ZT7.99±1.88 starting from the third day (Fig. 2E-F). In contrast, after 3 days of re-entrainment in 8L:16D, the OE-1 line showed a 3-hour phase delay (ZT 11.49±2.87) compared to Wt (Fig. 2E-F). Together, these results indicate that diatom cellular rhythmicity is entrained in a photoperiod-dependent manner and that *Pt*bHLH1a deregulation alters the capacity of cells to set diurnal phase pattern.

**Fig. 2.**
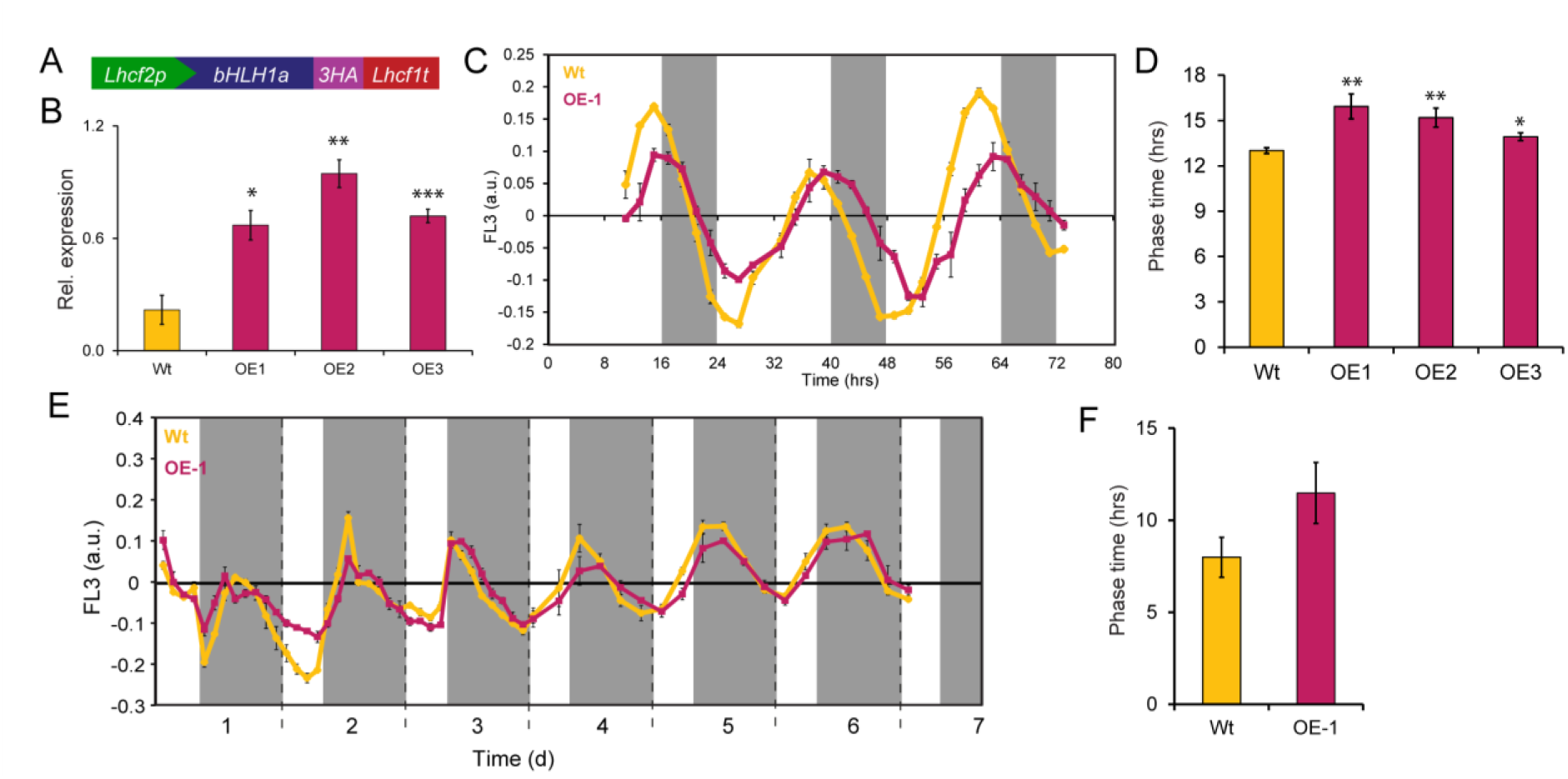
*PtbHLH1a* over-expression determines phase shifts in cellular rhythmicity. A) Schematic representation of the *Lhcf2p:bHLH1a-3xHA:Lhcf1t* construct used to generate *Pt*bHLH1a over-expressing lines. *Lhcf2p*: Light Harvesting Complex F2 promoter; 3HA: triple hemagglutinin tag; *Lhcf1t*: *Lhcf1* terminator. B) Quantification of total *PtbHLH1a* transcripts in OE-lines and Wt grown in 16L:8D photocycles and sampled at the ZT10. qRT-PCR expression values were normalized using *PtRPS* and *PtTBP* as reference genes and represent the average of 3 biological replicates (n=3) ± s.e.m (black bars). C) Diurnal oscillation of chlorophyll fluorescence (FL-3 parameter) in Wt and OE lines entrained under 16L:8D over three days. D) Diurnal phase time calculation of the FL-3 value in Wt and OE-1, OE-2 and OE-3 lines. Values represent the average of at least 4 biological replicates (n ≤ 4) from at least 2 independent experiments ± s.e.m. E) Phase re-entrainment analysis of fluorescence rhythms after photoperiod change from 16L:8D to 8L:16D in Wt (yellow) and OE-1 (red). The FL-3 parameter was monitored in cultures grown in 16L:8D photocycles and then transferred to 8L:16D for 6 days. Values represent the average of three biological replicates ± s.e.m. F) Bar plot representation of the re-entrained FL-3 phases in Wt and OE-1 cultures after three 8L:16D photocycles. Values represent the average of 3 biological replicates ± s.e.m. (black bars). *P<0.05, **P<0.01, ***P<0.001, t-test.

### *PtbHLH1a* regulates diurnal cell cycle progression

The altered rhythm of fluorescence upon *PtbHLH1a* over-expression described above may reflect delayed or asynchronous cell division dynamics. To get further insights into the effect of *PtbHLH1a* overexpression on rhythmic processes, cell cycle dynamics in the WT and OE-1 lines were thoroughly analysed. Cell cultures were synchronized by 40h of dark treatment and harvested on an hourly basis for 12h after re-illumination. At T0, total DNA content measurements showed comparable proportions of cells in G1 phase in all samples indicating effective synchronization of cell cultures (Fig. 3A). Starting after 3h of illumination, a progressive reduction of G1 cell number was observed in the Wt, with a minimum number reached after 10h of illumination. After 10h, the percentage of Wt cells in G1 increased with the emergence of daughter cells (Fig. 3A). Interestingly, compared to Wt, OE-1 cells showed a slower exit from the G1 phase, with the percentage G1 cells continuing to decrease over the entire 12h of illumination studied (Fig. 3A).

**Fig. 3.**
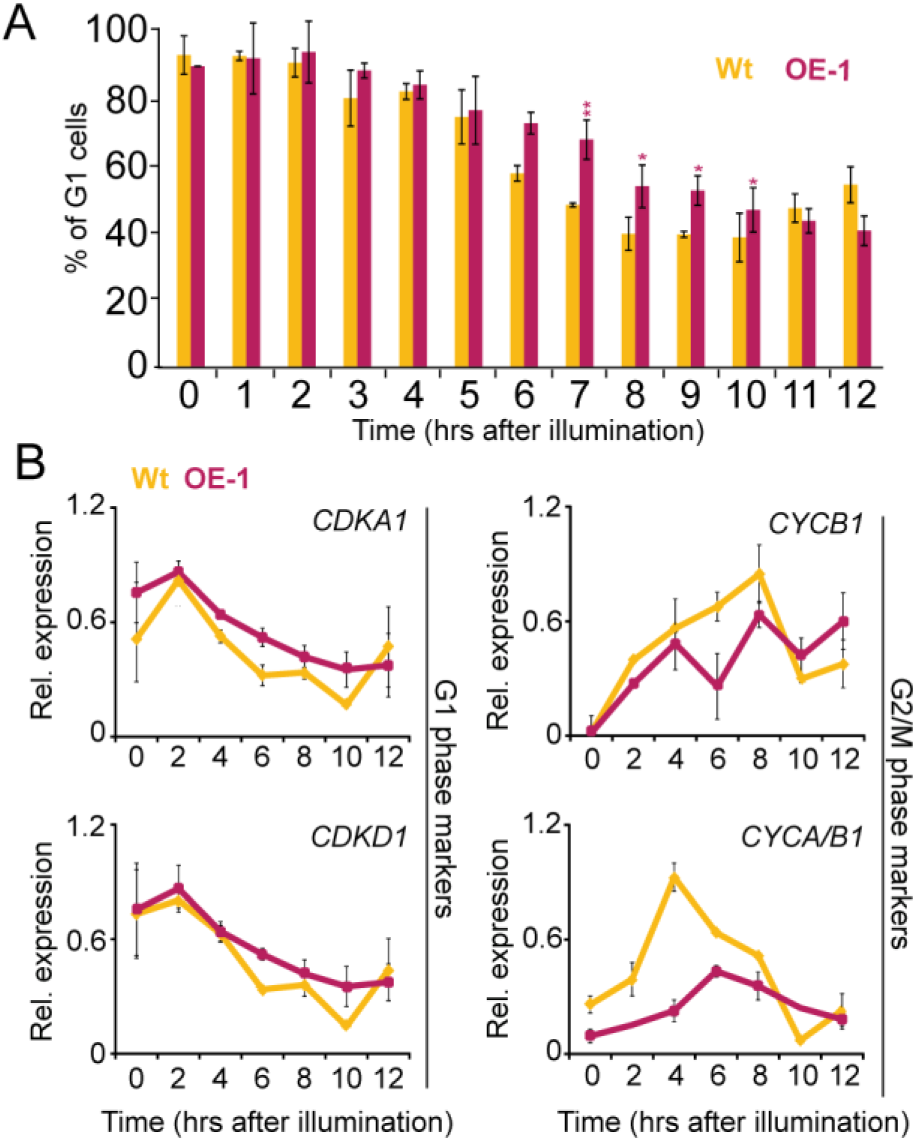
*PtbHLH1a* over-expression affects cell cycle progression. A) Cell cycle progression dynamics of Wt (yellow) and OE-1 (red) lines shown as the proportion of cells in the G1 phase measured by flow cytometry each hour over 12h of illumination following dark synchronization. Results are representative of three biological replicates ± s.e.m (black bars); t-test significance is indicated by *: P<0.05, **: P<0.01, t-test. B) qRT-PCR expression profiles of G1 (*PtCDKA1*, *PtCDKD1*) and G2/M phase marker genes (*PtCYCB1*, *PtCYCA/B1*) in synchronized Wt and OE-1 cell lines over 12h of illumination. Expression values were normalized using *PtRPS* and *PtTBP* as reference genes and represent the average of two independent biological replicates. For each gene, the expression value is relative to its maximum expression (maximum expression=1).

To further characterize the cell cycle deregulation caused by *PtbHLH1a* over-expression, the expression profiles of specific cell cycle phase marker genes (31) were analyzed in dark-synchronized Wt and OE-1 lines illuminated for 12h. The G1 phase gene markers *CDKA1* and *CDKD1* showed similar expression profiles in both lines until 4h from the onset of illumination (Fig. 3B). Starting from this time point, transcript levels of both genes were consistently higher in the OE-1 line compared to Wt except for at the end of the time course when they converged. This illustrates that the G1 phase duration of the two cell lines is different. Conversely, the G2/M marker *CYCB1* showed lower expression in OE-1 compared to the Wt between 4 and 8h after the onset of illumination (Fig. 3B). The expression of another G2/M phase marker, *CYCA/B1*, also resulted deregulated in OE-1, presenting reduced amplitude and peaking 2h later compared to Wt. Together, these results suggest that the deregulation of *PtbHLH1a* affects cell cycle, possibly by altering transition from G1 to S or G2/M phases.

### *Pt*bHLH1a regulates pace of diel gene expression

The effect of *Pt*bHLH1a de-regulation on gene expression was investigated since the expression of many *P. tricornutum* genes phase diurnally (27). To this end, Wt and OE-1 lines were grown in 16L:8D photocycles and sampled every 3 hours over 25 hours. For this analysis, genes with strong diurnal transcription oscillation were selected, including TFs (*bHLH1a*, *bHLH1b* and *bHLH3*) and rhythmic genes putatively involved in chlorophyll and carotenoid synthesis (*NADPH:protochlorophyllide oxidoreductase 2*, *Por2*, and *Violaxanthin de-epoxidase-related*, *Vdr*) (Fig. 1A, (27, 38)). Total *PtbHLH1a* transcript levels, including endogenous and transgenic mRNAs, were shown to be higher in OE-1 cells compared to the Wt, and the expression peaking at ZT7 in the OE-1 line and ZT10 in the Wt (Fig. 4A). A decrease of endogenous *PtbHLH1a* transcripts was observed in the OE-1 line compared to the Wt, possibly reflecting negative feed-back mechanism of *Pt*bHLH1a regulating its own transcription (Fig. 4A). A similar pattern was also observed for the *PtbHLH1b* gene, suggesting that *PtbHLH1a* and *PtbHLH1b* transcription is controlled by the same regulatory circuit. In addition, the *bHLH3* gene showed earlier phases of expression in OE-1 compared to the Wt (Fig. 4A). Similar deregulations of *PtbHLH1a, PtbHLH1b PtbHLH3* were also observed in the OE-2 and OE-3 lines at ZT10 (Fig. S3). Besides TFs, the Chlorophyll biosynthesis gene *Por2* was also anticipated and the *Vdr* gene presented increased amplitude of expression in the OE-1 line compared to the Wt (Fig. 4A).

**Fig. 4.**
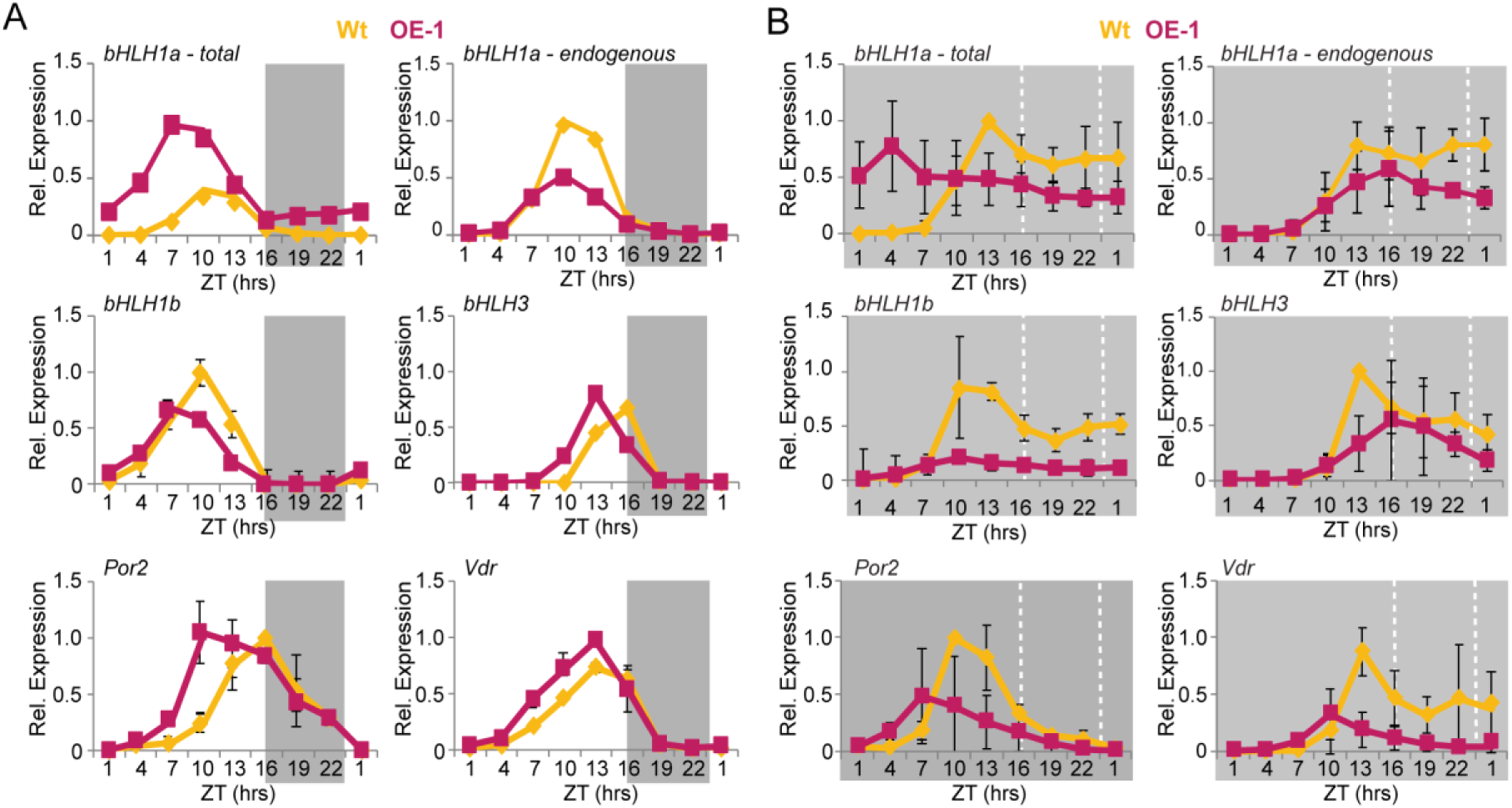
*PtbHLH1a* over-expression alters rhythmic diel gene expression. A) qRT-PCR diurnal expression analysis of *PtbHLH1a, PtbHLH1b*, *PtbHLH3*, *PtPor2* and *PtVdr* transcripts in 16L:8D entrained Wt (yellow) and OE-1 (red) cultures. *bHLH1a-total* represents the sum expression of the *bHLH1a* endogenous and the transgene transcripts; *PtbHLH1a-endogenous* refers to the endogenous gene only. Expression values represent the average of 2 biological replicates ± s.e.m. (black bars). B) nCounter expression analysis of *PtbHLH1a, PtbHLH1b*, *PtbHLH3*, *PtPor2* and *PtVdr* transcripts during 24 hours of dark free-running period in the Wt (yellow) and OE-1 (red) lines. Cells were previously entrained in 16L:8D cycles. Data represent the average of 3 biological replicates ± s.e.m (black bars). Grey rectangles represent dark periods. Expression values were normalized using *PtRPS* and *PtTBP* as reference genes. For each gene, the expression value is relative to its maximum expression (maximum expression=1).

Altered gene expression observed in *Pt*bHLH1a overexpression cell lines could be the consequence of the deregulation of cell cycle progression (Fig. 3). To test this hypothesis, gene expression was analyzed in dark conditions, when the cell cycle is arrested ((31) and (Fig. S5)). Because information about transcription dynamics in this condition was limited, an initial survey of expression of the previously selected 104 *P. tricornutum* diurnal rhythmic genes (Fig. 1A) was performed in cells exposed to continuous dark for 30 hours. Comparative analysis of transcript profiles revealed that around 20% of the genes show persistent oscillation of expression in D:D, although in some cases they displayed reduced amplitudes and/or shifted phases of expression compared to the 16L:8D condition. In particular, 19 genes were identified which showed the highest amplitude of expression in both L:D and D:D (for details see Materials and Methods), consisting of 16 putative TFs and 4 pigment-related enzymes (Fig. S6). Among the analyzed transcripts, genes that were severely affected by the absence of light were also found, being strongly down-regulated or over-expressed when compared to the L:D condition (Fig. S7). The expression of some of these genes was further analyzed in constant darkness in Wt and *Pt*bHLH1a OE-1 cells for a period of 24h. In the Wt, the analyzed genes showed comparable transcript profiles in D:D and 16L:8D conditions (Fig. 4B, Fig. S8). Conversely, 10 out of 13 tested genes displayed reduced amplitudes and shifts in the phase of expression in OE-1 compared to Wt in D:D (Fig. 4B, Fig. S8). It is worth mentioning that two of the analysed genes, *HSF1d* and *bZIP5*, showed almost overlapping profiles in OE-1 and Wt lines (Fig. S8), excluding global deregulation of transcription by modulation of bHLH1a expression. Taken together, these results suggest that *Pt*bHLH1a contributes to define timing of diurnal gene expression and that its activity is independent of direct light inputs and cell division.

### *Pt*bHLH1a-like proteins are widely represented in the genome of marine algae

bHLH-PAS proteins were thought to be restricted to the animal (Opistokonta) lineage (45) until genome and transcriptome sequencing projects revealed bHLH-PAS family members in diatoms (35) and other microalgae (46). Interestingly, when compared to animal bHLH-PAS, diatom proteins show peculiar features including a single predicted PAS domain (Fig. 5A), whereas animal bHLH-PAS proteins have two, and a N-ter extension that is absent in the animal counterparts. Available transcriptomic and genomic databases of marine algae and animals were searched for bHLH-PAS proteins and ≈90 novel bHLH-PAS proteins were discovered from Rhodophyta, Cryptophyta, Stramenopila, Alveolata and basal Opistokonta organisms (Table S3). With one exception, all the newly identified proteins showed a single predicted PAS domain, short C-ter extensions and N-ter regions of variable length, similar to the predicted structure of diatom bHLH-PAS proteins (Fig. 5A). Notably, the only bHLH-PAS possessing two PAS domains like the animal proteins was identified in *Galdieria sulphuraria* (Rhodophyta) and represents the first TF of this family identified in Archaeplastida. All the identified sequences, including selected bHLH-PAS from Opistokonta lineages, were used to perform a detailed phylogenetic analysis of the protein family using the bHLH and PAS domains. This analysis revealed at least three clades of algal bHLH-PAS proteins clearly separated from their Opistokonta counterparts (Fig. 5B). Interestingly, domain organization and branching positions of proteins from basal Opistokonta (*Monosiga brevicollis*) and microalgae (*Guillardia theta* (Cryptophyta) and *Nannochloropsis* (Stramenopila)) (Fig. 4B) support a possible common origin for this TF family, from an ancestor featuring single bHLH and PAS domains. However, the possible contribution of horizontal gene transfer and convergent evolution to the proliferation and diversification of this family cannot be excluded, and may be supported by the features of the *G. sulphuraria* bHLH-PAS protein, likely independently acquired by this alga. Based on our analysis, the majority of microalgal bHLH-PAS proteins fall into three separate clades: the first containing 9 TFs from diatoms and *Ectocarpus siliculosus*, the second comprising *Pt*bHLH1a together with 35 proteins from diatoms and Alveolata (Dinoflagellata), and the third comprising 41 proteins from Alveolata (Ciliophora and Dinoflagellata) and diatoms, including *Pt*bHLH1b (Fig. 5B).

**Fig. 5.**
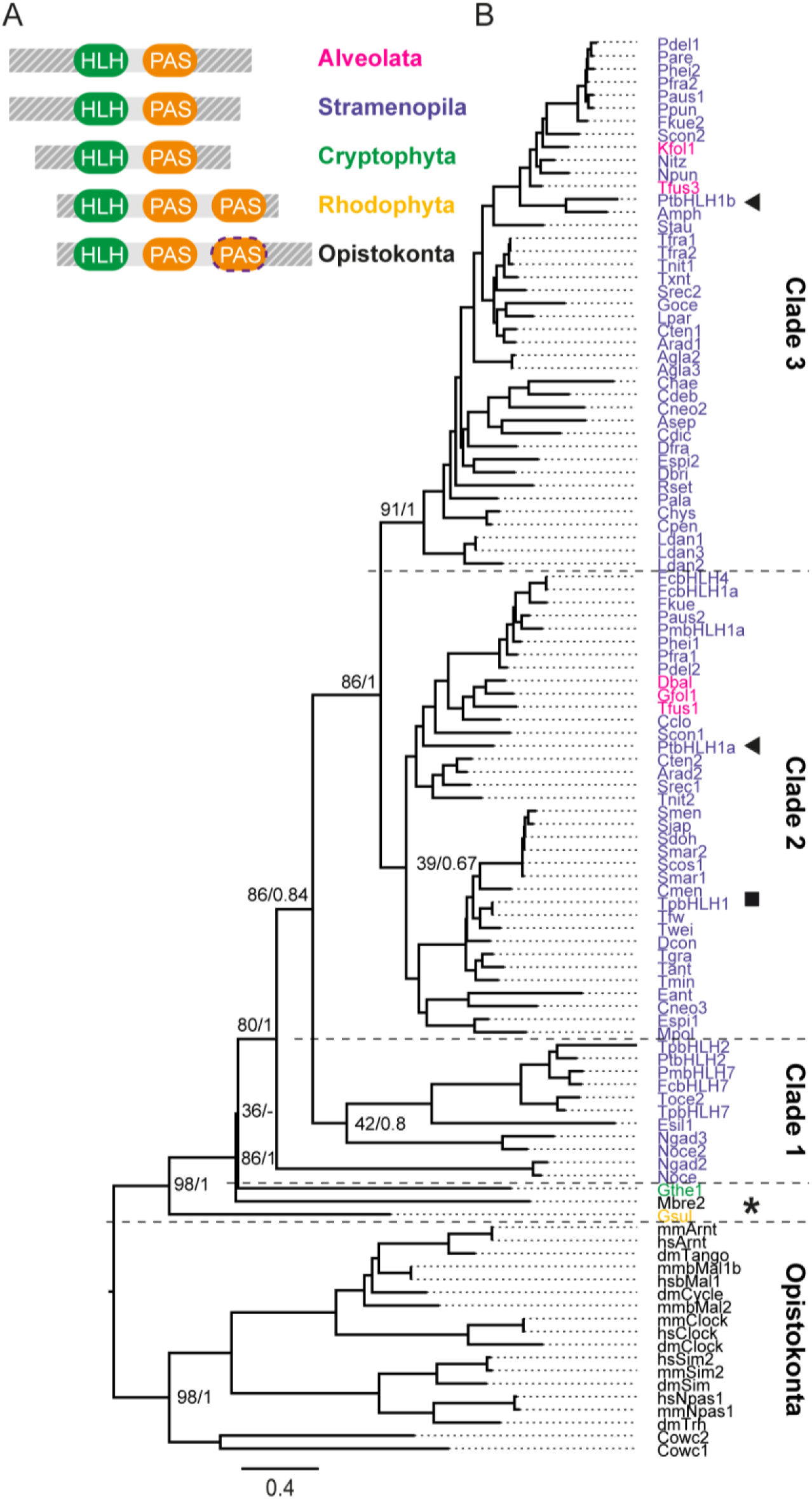
bHLH-PAS protein family structure and phylogeny. A) Schematic representation of bHLH-PAS protein domain architecture across Eukaryotes. Segmented line indicates possible absence of the second PAS domain in some Opistokonta species. The grey patterns before the bHLH domain and after the PAS domain represent the variations in N-ter and C-ter length in different organisms. B) Maximum Likelihood (ML) phylogenetic tree of the bHLH-PAS family. The Opistokonta clade is used as the outgroup. Numbers refer to bootstrap values of the basal nodes using ML (RAxML, 1000 bootstraps) and Bayesian Inference (MrBayes, 2.5M generations, 25% burn-in). The asterisk, the arrows and the square indicate the position of *Monosiga brevicollis* bHLH-PAS, *P. tricornutum* HLH1a and bHLH1b, and *Thalassiosira pseudonana* bHLH1, respectively. The colour code indicates the lineage corresponding to each bHLH-PAS protein, shown in Fig 5A.

Our results highlight diversification and widespread distribution of bHLH-PAS family members in different groups of algae. Moreover, the presence of bHLH1a-like genes in the genome of dinoflagellate and diatom species suggests that these proteins may share similar functions in microalgae. The similarities in diel transcript regulation and timing of expression between *Pt*bHLH1a and the centric diatom *Thalassiosira pseudonana* ortholog, *TpHLH1* (26), further reinforce this hypothesis.

## DISCUSSION

The diurnal cycle is characterised by profound periodic light and temperature changes which have shaped the evolution of most ecosystems on Earth. In most organisms, biological rhythms are controlled by interconnected transcriptional-translational feed-back loops involving TFs and integrating signals from the environment (5). Although this regulatory framework is conserved among eukaryotes, the regulators responsible for the timing of events within biological rhythms seem to have emerged several times through evolution (47). Therefore, our current understanding of diurnal and circadian regulation, largely based on the study of terrestrial model organisms, is not always appropriate or relevant for evolutionarily distant marine organisms. In this study, we have shed light on the unknown regulators of diurnal patterns in diatoms, one of the most prominent phytoplanktonic groups in the Ocean. In agreement with previous studies (27, 28), we showed the existence of organized transcriptional programs defining cellular activities along the daily cycle in *P. tricornutum* and identified a number of TFs phasing at different times during the 24h cycle, as novel candidates for diatom diurnal regulation. By monitoring diurnal variations in chlorophyll fluorescence, robust regulation of diatom physiological rhythms that can be re-entrained to changing photoperiods was also unveiled. The ability to adjust the phase according to the photoperiod length constitutes one defining criterion of circadian clock-regulated mechanisms (48). Likewise, the strongly oscillating diel expression pattern of a *P. tricornutum* bHLH-PAS gene, *PtbHLH1a*, responded to photoperiod length with peak expression 4 hours before night onset in both 12h and 16h day photoperiods. The timing of *PtbHLH1a* expression is also preserved in cells under iron deficiency, in contrast to the expression of many other *P. tricornutum* genes observed previously, and despite the growth rate reductions caused by nutriment depletion (28). Functional characterization of *Pt*bHLH1a established its involvement in the regulation of *P. tricornutum* diurnal rhythms. Transgenic lines over-expressing *PtbHLH1a* using a promotor that is activated earlier in the light period than the endogenous gene maintained cellular rhythms of ∼24 h but show phase-alterations that are even more accentuated in re-entrainment experiments. This phenotype may reflect a reduced capacity of cells to synchronize to environmental light-dark cycles and adjust the phase to the new photoperiod. The participation of bHLH1a in the regulation of *P. tricornutum* cell cycle progression, reported in this study, could also explain the altered cellular fluorescence rhythmicity observed in the mutants. Altered cell division timing could derive from a delayed exit from the G1 phase in transgenic lines compared to Wt, as also supported by the altered expression of the mitotic cyclins CYCB1 and CYCA/B1 in these lines (31). The *Pt*bHLH1a gene could participate in gating cell divisions at night time, therefore maximizing the energetic budget, as observed in several unicellular algae (7, 8, 30, 49). Interestingly, a similar regulation of the cell cycle occurs in mammalian cells, where the circadian clock controls the expression of G2 cycle-related genes to gate cell division at specific times of the day (50).

Besides cell cycle, *Pt*bHLH1a deregulation also affected diurnal rhythmicity of several gene transcripts. This phenotype was uncoupled from cell cycle deregulation as it was observed also in the absence of cell division, during darkness. Interestingly, the deregulation of gene transcription was much more pronounced when analyzed in D:D compared to L:D conditions. These results suggest on one hand that multiple regulatory inputs participate in the regulation of diurnal rhythmic gene transcription (48), partially masking *Pt*bHLH1a contribution to this process in cyclic environments, and, on the other hand, support *Pt*bHLH1a involvement in the maintenance of rhythms in the absence of light inputs.

The evidence provided in this work support the hypothesis that *Pt*bHLH1a is one component of an uncharacterized endogenous circadian clock in diatoms, either as part of a central oscillator or as a mediator of clock inputs or outputs. With the exception of CRY (33, 39), orthologs of plant and animal circadian clock genes are absent in diatom genomes. However, *Pt*bHLH1a contains bHLH and PAS protein domains that are also present in the CLOCK and BMAL proteins, components of the mammalian central circadian oscillator (51, 52). Interestingly, previous studies showed that the *P. tricornutum* animal-like blue light sensor Cpf1 can repress the transcriptional activity of these proteins in a heterologous mammalian cell system (33), suggesting at least partial conservation in the regulatory program generating rhythmicity in animals and diatoms. The downregulation of endogenous *PtbHLH1a* and *PtbHLH1*_*b*_ transcripts in the OE lines analysed in this work likely reflects a negative feedback loop in the regulation of these genes, which is typical of circadian genetic oscillators, and further supports a possible role for *PtbHLH1a* in diatom rhythm regulation. Moreover, as observed for the clock components, we show that *Pt*bHLH1a contributes to set the phase of output processes in cycling environments and to the maintenance of gene expression rhythmicity in constant darkness. Some TFs characterized in this study and showing altered expression patterns in *Pt*bHLH1a transgenic cells (*i.e., bHLH1b*, *bHLH3*, *bZIP5*, *bZIP7*, *HSF1d*, *HSF1g*, *HSF3.3a*, *HSF4.7b*) represent direct or indirect targets of *Pt*bHLH1a activity and possible additional components of the network participating in diel rhythm regulation. *Pt*bHLH1a might act downstream of signal transduction cascades activated by the diatom photoreceptors analysed in this study and elsewhere. The presence of a PAS domain in *Pt*bHLH1a also suggests that this protein might have its own light-sensing ability (53).

Although further analyses under prolonged free running conditions (for example continuous darkness and continuous illumination) will be necessary to conclusively assess the involvement of *Pt*bHLH1a in circadian regulation, this protein constitutes a promising entry point for the characterization of diatom molecular timekeepers. Finally, the discovery of the wide distribution of bHLH-PAS domain-containing proteins in diatoms, as well as in other algae, has the potential to shed new light on the evolution of biological rhythms. bHLH-PAS proteins might have independently acquired their function in rhythm regulation by convergent evolution. However, the existence of this function in an ancient heterotrophic marine ancestor that subsequently acquired plastids via endosymbiosis events (54) and prior to colonization of land cannot be excluded. Regulators of cellular rhythmicity such as *Pt*bHLH1a may have played a critical role for diatom prominence in marine ecosystems, by synchronizing cellular activities in optimal temporal programs and maximizing diatoms’ ability to anticipate and adapt to cyclic environmental variations.

Biological rhythms are still poorly understood at molecular and mechanistic levels in marine algae, despite their fundamental significance to these organisms’ biology and ecology. Further characterization of *Pt*bHLH1a homologs in diatoms and other algae is expected to provide new insights into biological rhythms in marine organisms.

## METHODS

### Culture conditions

Wild-type *P. tricornutum* (*Pt*1 8.6; CCMP2561) cells and transgenic lines were grown at 18°C with shaking at 100 rpm in F/2 medium (55) without silica and illuminated at 40 µmol photons m^-2^ s^-1^ of white light (Philips TL-D De Luxe Pro 950). Detailed information is in SI.

### Cell cycle analysis

Cells were synchronized in the G1 phase by 40h of darkness. After re-illumination, samples were collected every hour for 12h. Details are in SI.

### RNA extraction and gene expression analyses

Total RNA was extracted and qRT-PCR performed as described in (36). Codeset information and raw nCounter data are available from the GEO database (<u>Series GSE112268</u>). Detailed information is in SI.

### Selection of rhythmic transcripts and clustering analysis

To select genes with rhythmic expression in the light-dark cycle, we used microarray data from (27). To select rhythmic transcripts in D:D, standard deviation from the average expression were calculated and used as selective criteria. Detailed information is in SI.

### Generation of the *PtbHLH1a* overexpressing lines

Diatom transgenic lines were obtained by co-transformation of the pDEST-C-HA-*PtbHLH1a* plasmid together with Nourseothricin resistance plasmid. Details are described in SI.

### Data mining, protein sequence and phylogenetic analysis

Detailed information about data mining, protein sequence and phylogenetic analysis is provided in SI.

## ACKNOWLEDGEMENTS

We thank M. Jaubert and L. De Veylder for critical suggestions, A. Manzotti for help with characterization of OE lines, P. Oliveri for support with nCounter technology, D. Petroutsos and G. Finazzi for support in monitoring cell physiology. This work was funded by the HFSP research grant (#RGY0082/2010), EMBRIC, EMBRC-FR and a grant from the Gordon and Betty Moore Foundation (GBMF 4966) to A.F.

## SUPPORTING INFORMATION

### Culture conditions

For experiments in different photoperiods, cultures were pre-adapted to the different L:D cycles for 2 weeks before starting the experiment. For experiments in continuous darkness cells were pre-adapted in 16L:8D photocycles for 2 weeks, then transferred to D:D at the start of the experiment. Growth measurements were performed using a MACSQuant Analyser flow cytometer (Miltenyi Biotec, Germany) by counting the cells based on the R1-A (630nm excitation, 670-700nm emission) versus the R1-H parameters. Phase rhythmicity assays were carried out by measuring the Chlorophyll fluorescence using the flow cytometer FL-3 parameter (488 nm excitation, 670-700 nm emission). For the re-entrainment experiment, cultures were initially entrained under 16L:8D photocycles at 40 µmol m^-2^ s^-1^ of white light, then transferred to 8L:16D photocycles at 80 µmol m^-2^ s^-1^ of white light for 6 days. All phase time and period calculations were performed using the MFourfit curve-fitting method using the Biodare2 tool (biodare2.ed.ac.uk, (1)).

### Cell cycle analysis

For cell cycle analysis cells were pelleted by centrifugation (4000 rpm, 15 minutes, 4°C), fixed in 70% EtOH and stored in the dark at 4°C until processing. Fixed cells were then washed three times with 1×PBS, stained with 4’,6-diamidino-2-phenylindole (at a final concentration of 1 ng/ml) on ice for 30’, then washed and resuspended in 1xPBS. After staining, samples were immediately analyzed with a MACsQuant Analyser flow cytometer (Miltenyi Biotec, Germany). For each sample, 30,000 cells were analysed and G1 and G2 proportions were inferred by calculating the 2c and 4c peak areas at 450 nm (V1-A channel) using the R software. A peak calling method was applied to the resulting histogram, based on a 1^st^ derivative approach (2). The locations of G1 and G2 peaks were first determined using G1 and G2 reference samples and then used to identify G1 and G2 cells in the experimental samples. The area under each peak was used as a proxy for the proportion of cells in each population.

### RNA extraction and gene expression analyses

For qRT-PCR analysis *PtRPS* and *PtTBP* were used as reference genes. Each independent replica of the qRT-PCR data was normalized against the maximum expression value of each gene (*i.e.,* gene expression range lies between 0 and 1 across the time series). Average expression and standard error was then calculated and plotted. The full list of oligonucleotides used in this work can be found in Table S4. For the nCounter analysis, gene specific probes (Table S1) were designed and screened against the *P. tricornutum* annotated transcript database (JGI, genome version 2, Phatr2) for potential cross-hybridization. Total RNA extracts (100 ng) from three biological replicates were used for hybridization. Transcript levels were measured using the nCounter analysis system (Nanostring Technologies) at the UCL Nanostring Facility (London, UK) and at the Institut Curie technical platform (Paris, France) as previously described (3). Expression values were first normalized against the internal spike-in controls, then against the geometric mean of the 3 reference genes *PtRPS*, *PtTBP* and *PtACTIN12*.

### Selection of rhythmic transcripts and clustering analysis

For the selection of genes with rhythmic expression in the light-dark cycle, we used microarray data from (4). First, we identified all the genes belonging to the TFs, photoreceptors, cell cycle and metabolism-related categories (pigment synthesis and photosynthesis). Then, transcripts were ranked based on a defined t-value for each time point (mean gene expression of the replica/(1+s.d.)) and those showing t-value >+0.7 or <-0.7 across the time series, were retained. The nCounter expression data was normalized against the maximum expression value of each gene, in a similar way to qRT-PCR expression data. This normalization was applied to the 3 replicas independently and for each condition (L:D and D:D) with the average expression value used for the clustering analysis. Hierarchical clustering analysis was performed with MeV 4.9 (5) using Pearson correlation. Peak analysis was performed using the MFourfit curve-fitting method defining average expression phases for each cluster (biodare2.ed.ac.uk, (1)).

For the selection of rhythmic transcripts in D:D, expression values were normalized using as reference genes *PtRPS, PtTBP* and *PtACTIN12* and the genes with the highest values of standard deviation from the average expression (M value) over the two time courses (16L:8D and D:D) were selected. A threshold equal to 1 was set using the published *P. tricornutum* diurnal microarray dataset (4) as background. Gene expression profiles were further empirically examined and false positives eliminated.

### Generation of the *PtbHLH1a* overexpressing lines

Transformed cells were tested for the presence of the transgene by PCR and qRT-PCR analysis (see Table S4 for oligonucleotide sequences). The full length *PtbHLH1a* coding sequence was obtained by PCR amplification with the specific oligonucleotides *Pt*bHLH1a-DraI-Fw and *Pt*bHLH1a-XhoI-Rv on cDNA template using the Phusion high fidelity DNA polymerase (Thermo Fisher, USA). The PCR fragment was inserted into the pENTR1A vector (Invitrogen, USA) using the DraI/XhoI restriction sites, and recombined with the pDEST-C-HA vector (6).

### Data mining, protein sequence and phylogenetic analysis

The *P. tricornutum* bHLH1a (Phatr3_J44962) protein sequence was used as the query for blastP analyses on the JGI, NCBI and MMETSP public database (7). Searches of the pfam database for proteins possessing both the HLH and PAS domains were also performed. The retrieved sequences were analyzed using the batch search tool on the CDD (Conserved Domain Database) NCBI server to retrieve proteins presenting at least one HLH and one PAS domain only. We identified 100 HLH-PAS proteins from 71 marine algal species which were aligned using MAFFT (8), along with 22 HLH-PAS proteins from relevant metazoan (*Homo sapiens*, *Mus musculus* and *Drosophila melanogaster*) and unicellular Opistokonta (*Monosiga brevicollis* and *Capsaspora owczarzaki*). Preliminary phylogenies were produced with MEGA 7 (9) to eliminate ambiguously aligned sequences, refining the alignment to 107 sequences and a final length of 198 aa (<5% gap per position). The best aminoacidic model to fit the data was estimated with ProtTest 3.4.2 (10). Phylogenetic analyses were performed with RAxML (1000 bootstraps) and MrBayes 3.2.6 (2.5 million generations, 2 runs, 25% burn-in) on the CIPRESS gateway (11). The final tree was edited in FigureTree 1.4 (http://tree.bio.ed.ac.uk/software/Figuretree/). GenBank accession codes of the genes utilized in the bHLH-PAS phylogenetic analysis are reported in Table S3.

---

**Fig. S1.**
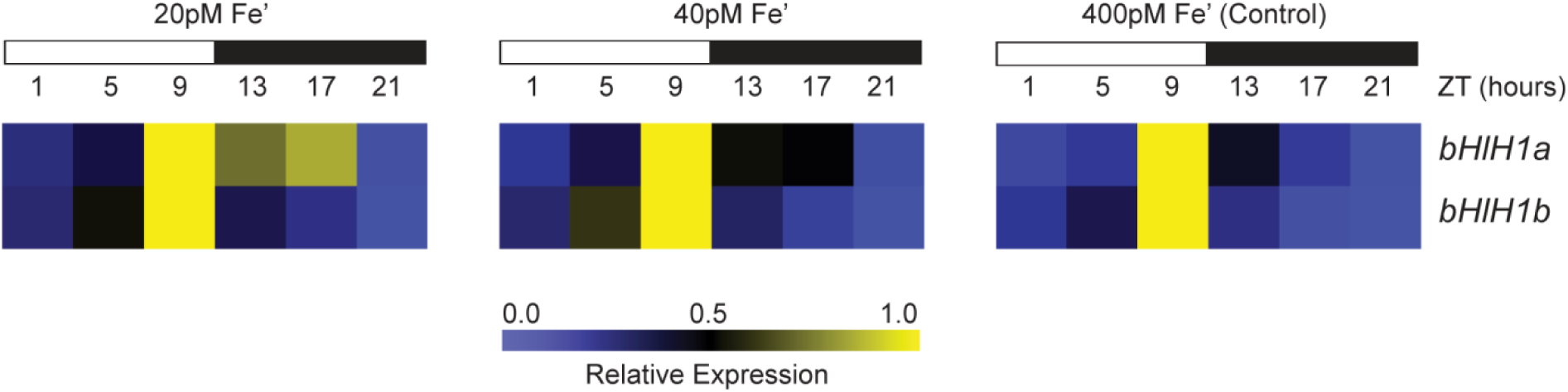
Diel expression patterns of *PtbHLH-PAS* genes under Fe-depletion conditions in 12L:12D photoperiods. Diel expression patterns of *PtbHLH1a* and *bHLH1b* in normal (400 pM Fe’) and iron depletion conditions (40 and 20 pM Fe’) were obtained using transcriptome data extracted from (36). Light and dark periods are represented by white and black rectangles. Expression values are given relative to the maximum expression for each gene, where ‘1’ represents the highest expression value of the time series.

**Fig. S2.**
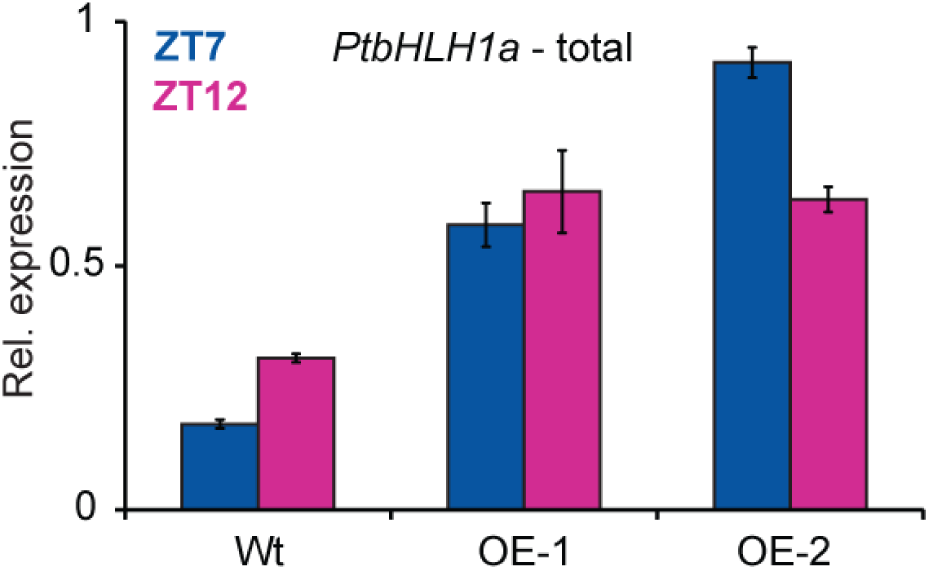
The *Lhcf2p:bHLH1a-3xHA:Lhcf1t* construct drives over-expression and anticipation of *PtbHLH1a*. Quantification of total *PtbHLH1a* transcripts in two independent *Pt*bHLH1a over-expressing lines (OE-1, OE-2) compared to the wild type (Wt) strain. Cells were grown in 16L:8D photocycles and sampled at the ZT7 and ZT12 time points. qRT-PCR expression values were normalized using *PtRPS* and *PtTBP* as reference genes and represent the average of two biological replicates (n=2) ± s.e.m (black bars).

**Fig. S3.**
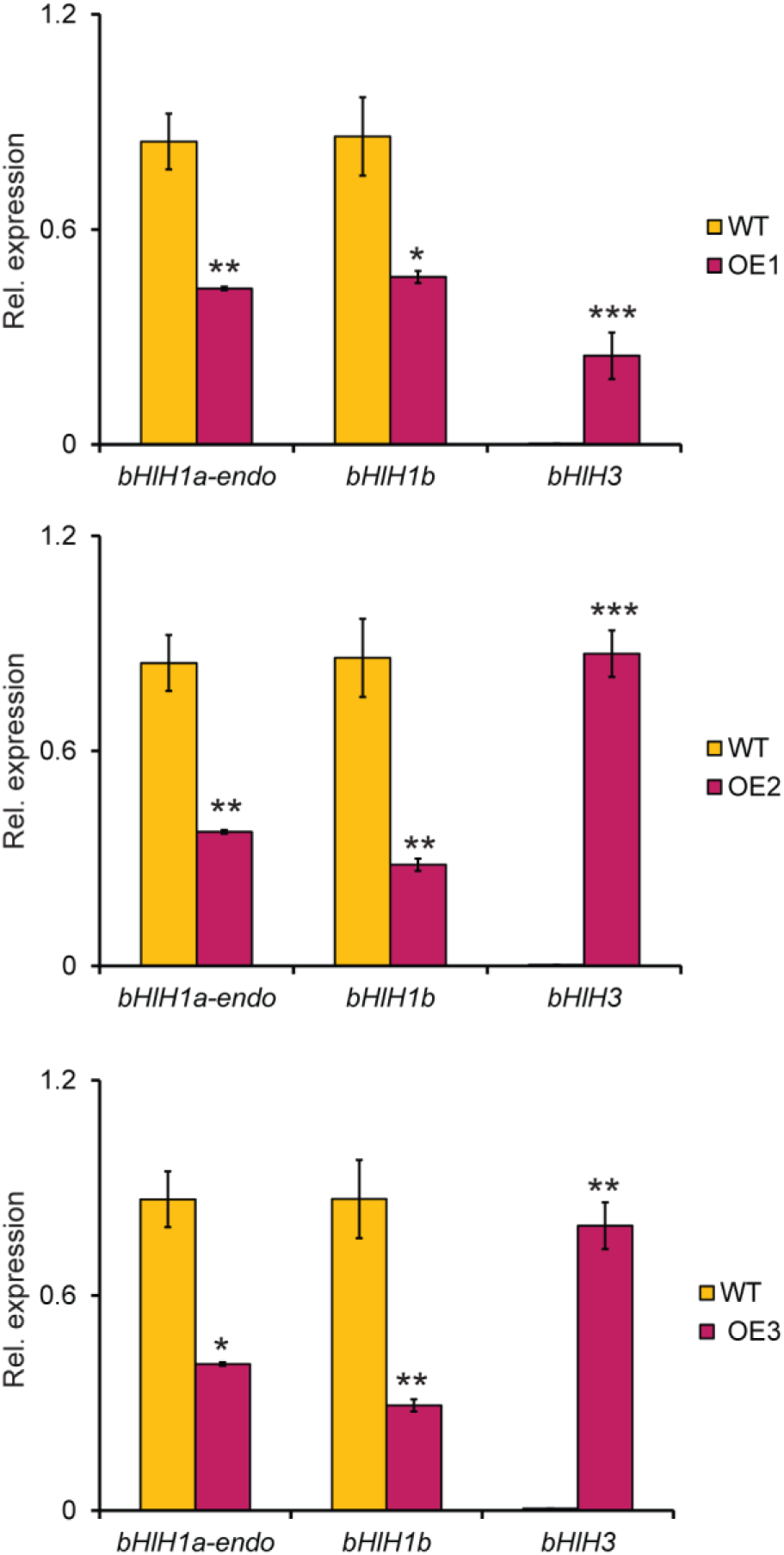
*PtbHLH1a* over-expression alters rhythmic diel gene expression in three independent lines. Quantitative gene expression analysis by qRT-PCR of *PtbHLH1a-endogenous, PtbHLH1b* and *PtbHLH3*, transcripts in 16L:8D entrained Wt, OE-1, OE-2 and OE-3 cultures. Samples were harvested at ZT10. *PtbHLH1a-endo* refers to the transcript levels of the endogenous gene only. Expression values represent the average of 3 biological replicates ± s.e.m. (black bars) and were normalized using *PtRPS* and *PtTBP* as reference genes. For each gene, expression values are represented as relative to its maximum expression corresponding in the graph to ‘1’. *P<0.05, **P<0.01, ***P<0.001, t-test.

**Fig. S4.**
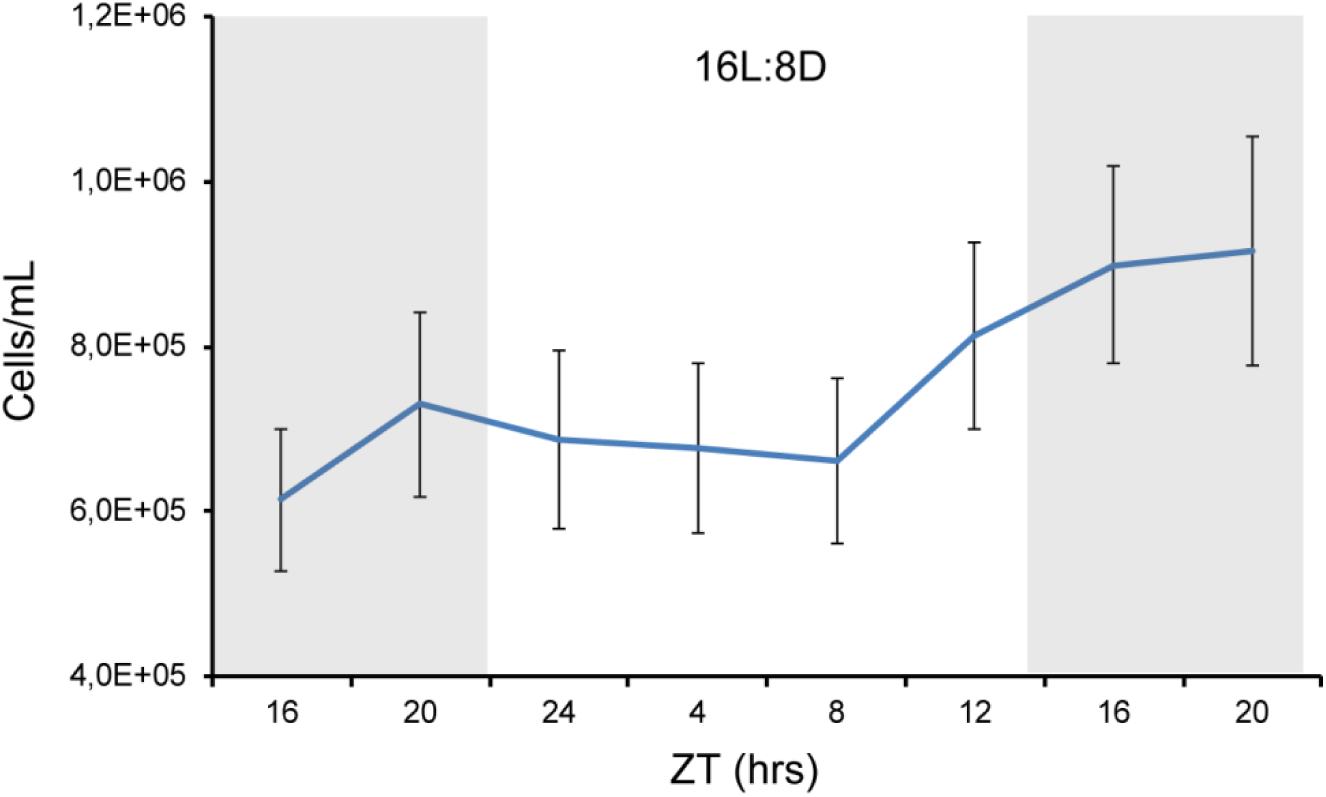
Diurnal cell growth dynamics in *P. tricornutum.*. Diel cell number measurements in wild type cultures grown under 16L:8D photoperiods. Values represent the mean counts of three independent biological replicates ± s.e.m. (black bars).

**Fig. S5.**
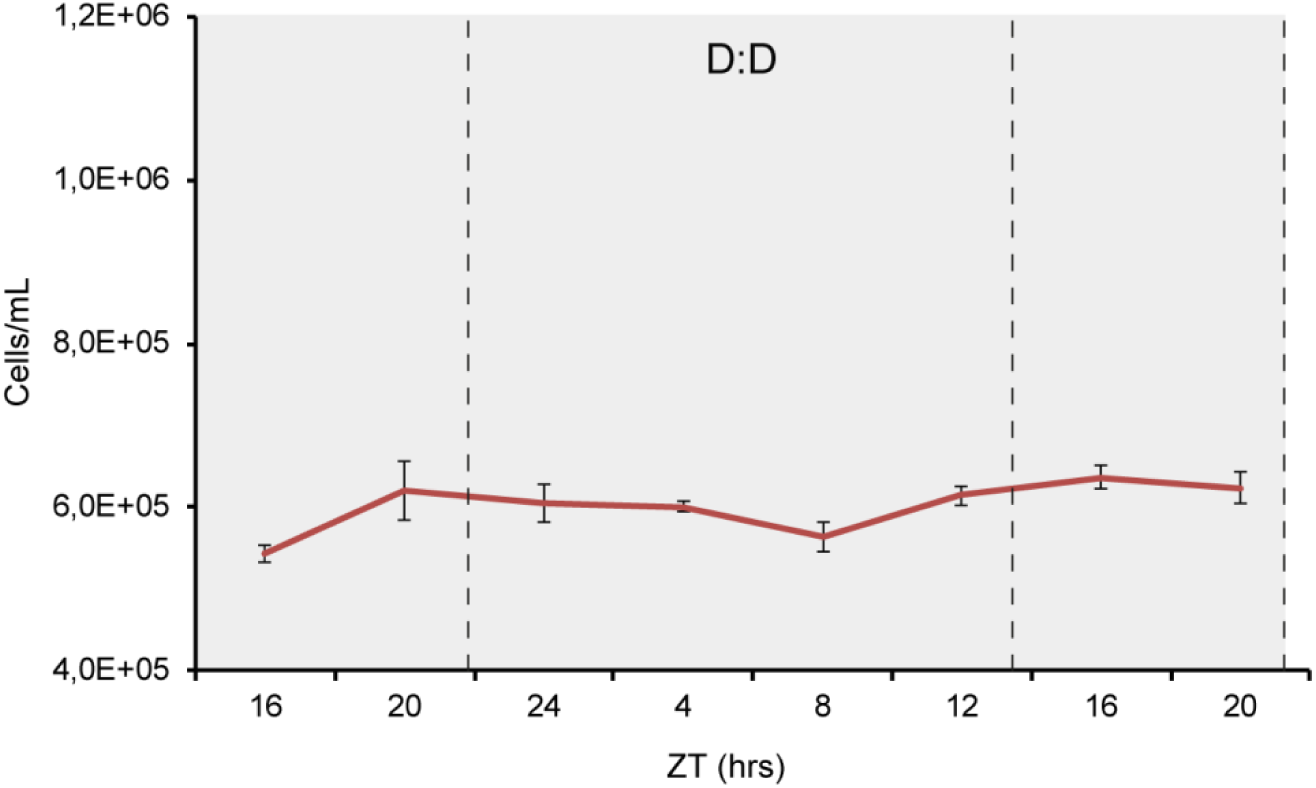
Cell division is arrested in continuous dark conditions in *P. tricornutum*. Diurnal cell growth measurements of the wild type cultures entrained in 16L:8D and then transferred to continuous darkness condition (D:D). Values represent the mean counts of three independent experiments ± s.e.m. (black bars).

**Fig. S6.**
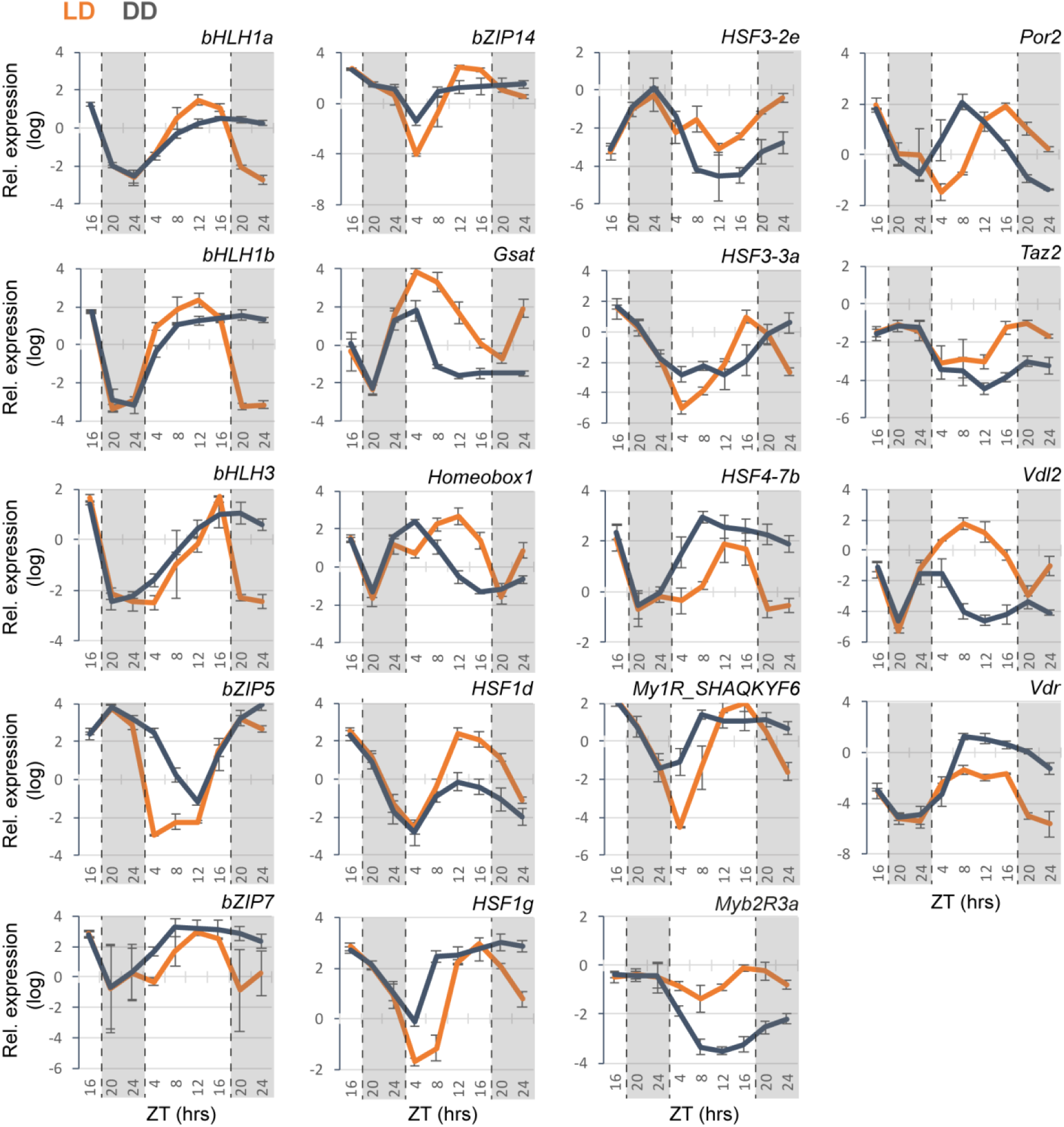
nCounter expression analysis of genes maintaining rhythmic expression in D:D conditions and 16L:8D condition in Wt cells. Data represent the average expression of biological triplicates ±SD and are normalized using the *PtRPS*, *PtTBP* and *PtACTIN12* reference genes. For each gene, the expression value is relative to its maximum expression corresponding in the graph to ‘1’. Results for cells grown in 16L:8D cycle are shown in orange (L:D); Results for cells in constant darkness (following 16L:8D adaptation) are shown in grey (D:D).

**Fig. S7.**
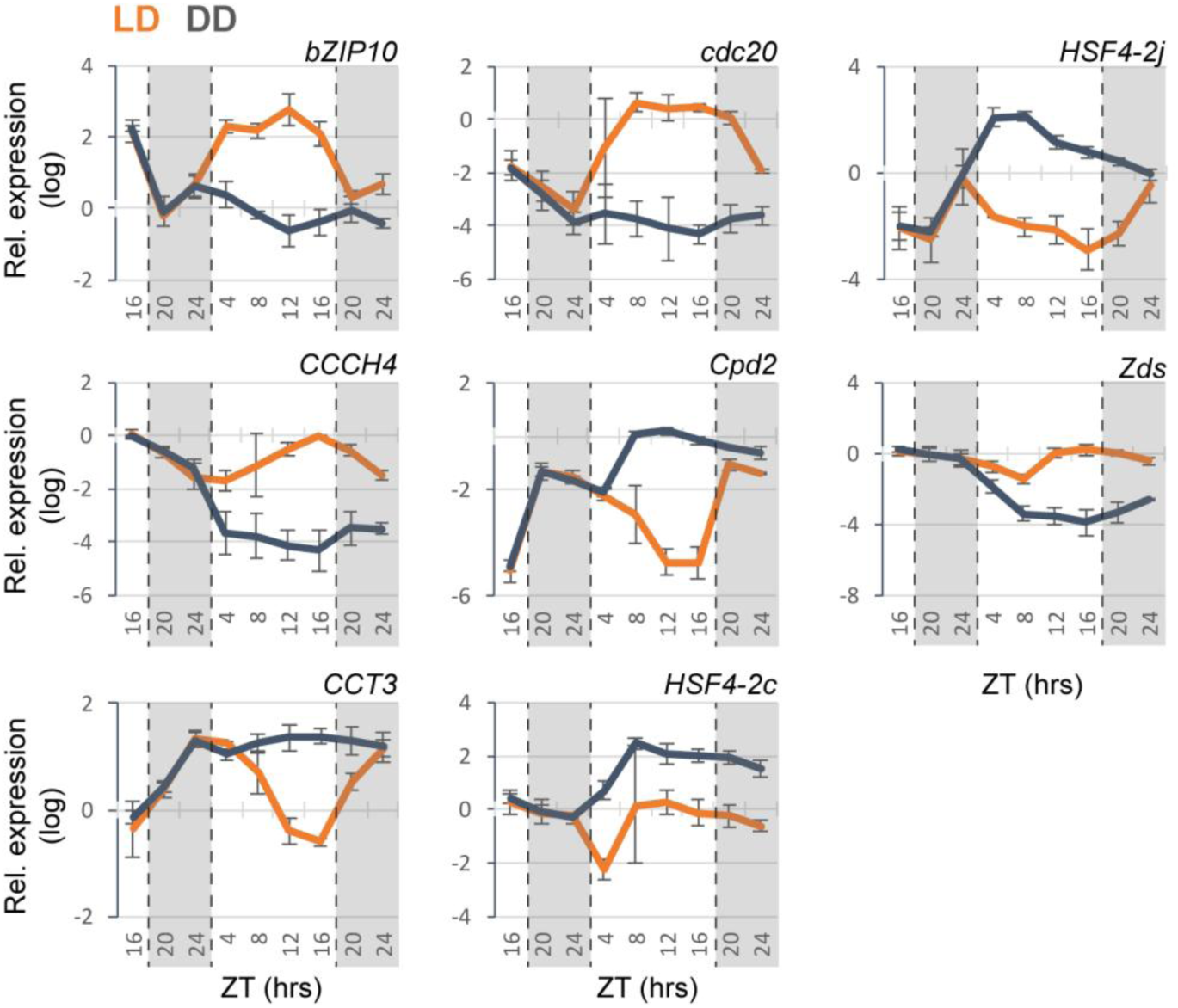
nCounter expression analysis of selected genes with altered rhythmic expression in Wt cells in D:D conditions compared to 16L:8D condition. Expression values represent the average of three biological triplicates ±SD and are normalized using the *PtRPS*, *PtTBP* and *PtACTIN12* reference genes. For each gene, the expression value is relative to its maximum expression corresponding in the graph to ‘1’. Results for cells grown in 16L:8D cycle are shown in orange (L:D); Results for cells in constant darkness (following 16L:8D adaptation) are shown in grey (D:D).

**Fig. S8.**
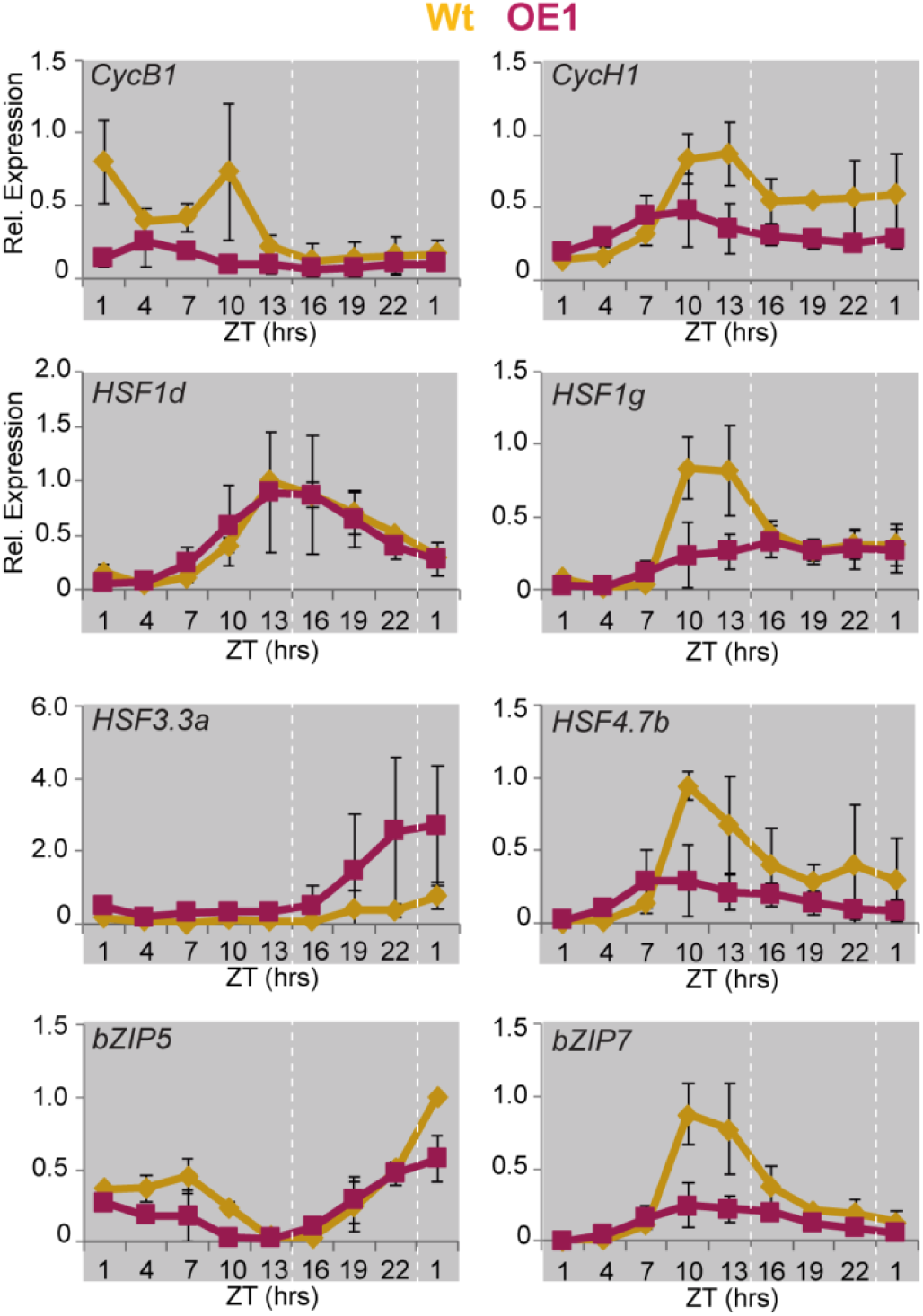
nCounter analysis of selected rhythmic gene expression profiles in continuous darkness in Wt and *Pt*bHLH1a OE-1 cells. Cells were entrained in 16L:8D cycles, then transferred to D:D and collected every 3 hours for 24h. Expression values represent the average of three biological replicates (n=3) ± s.e.m (black bars) and have been normalized using *PtRPS* and *PtTBP*.

**Table S1:**
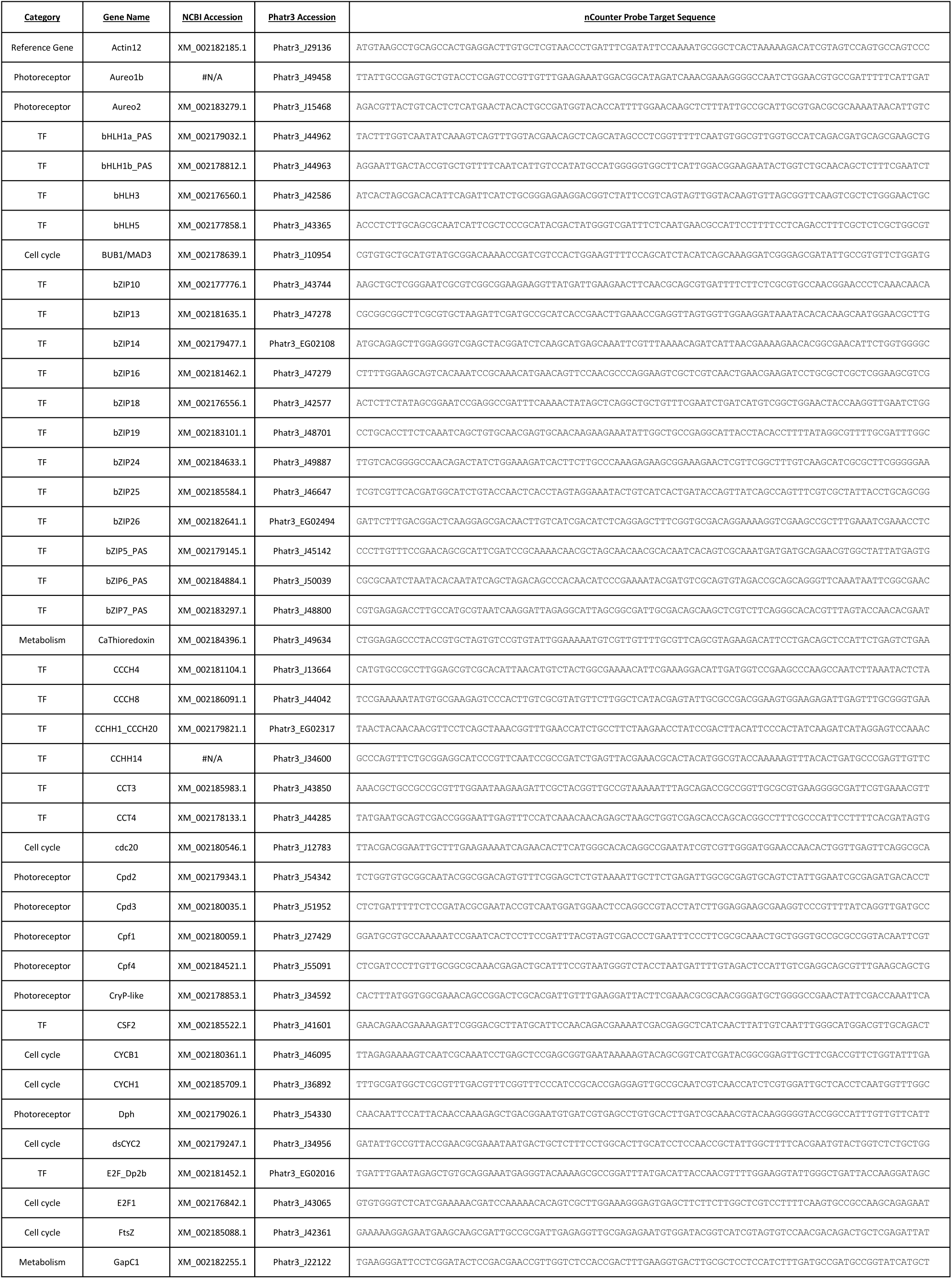
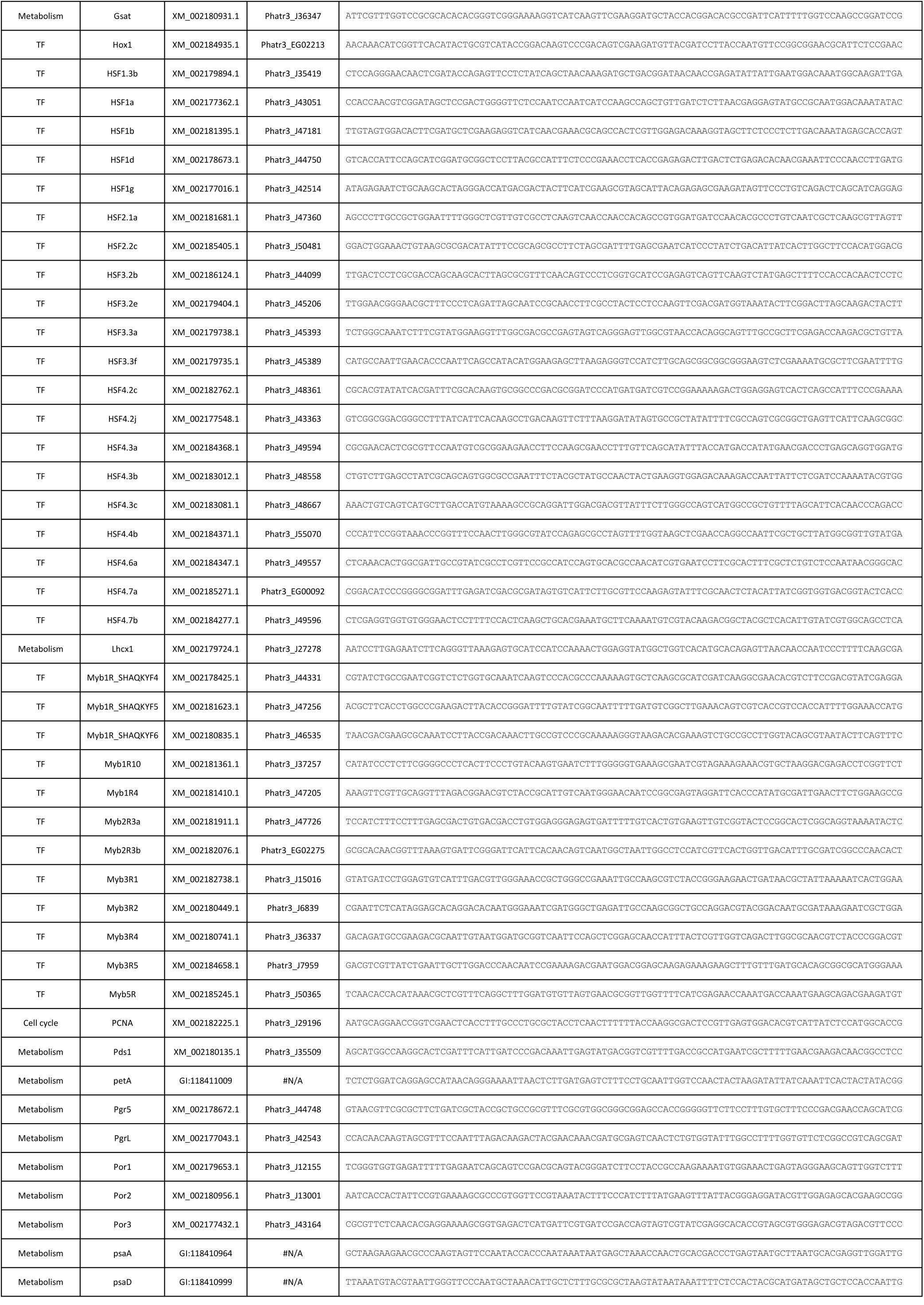
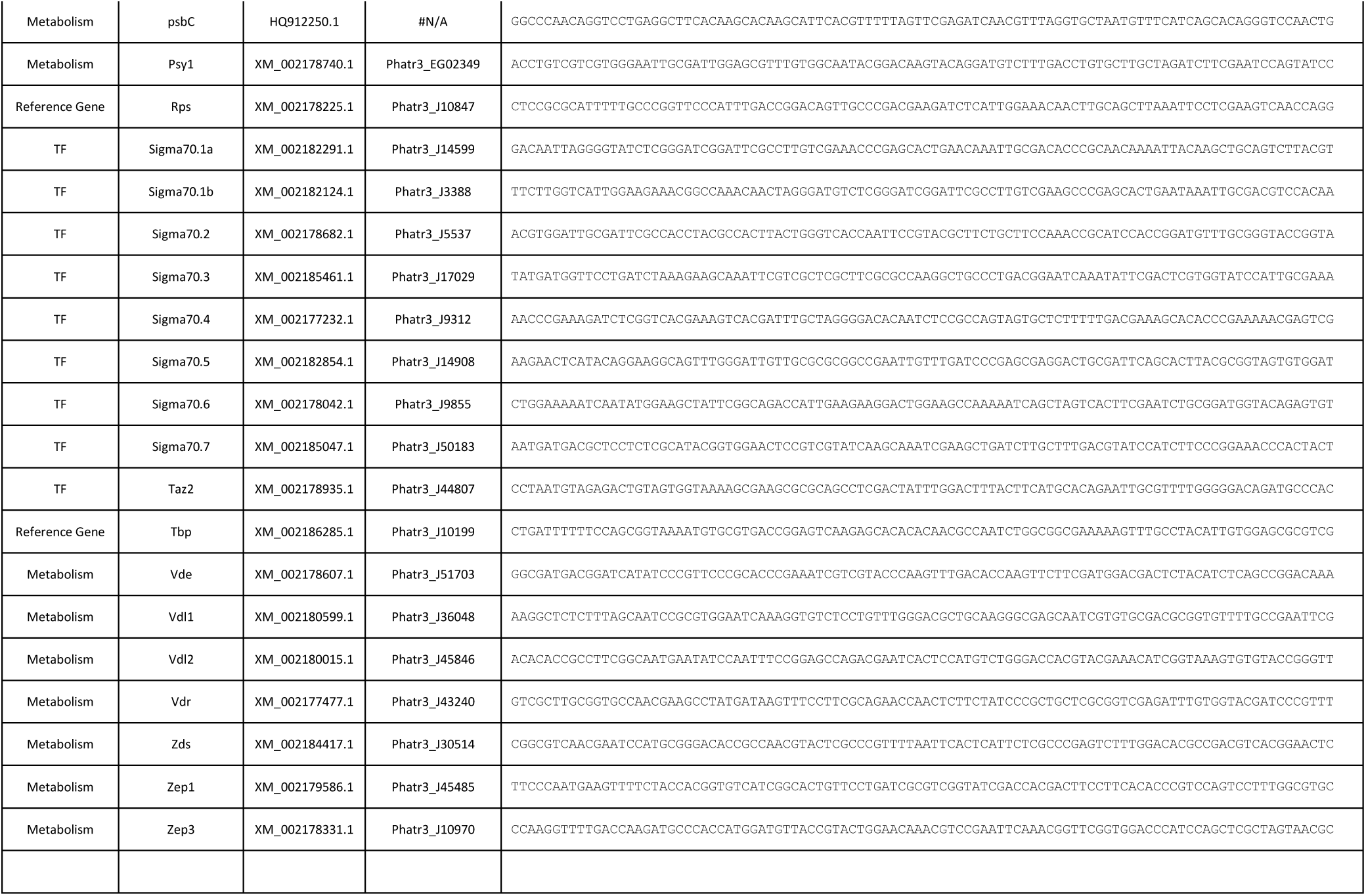
Accession codes and sequence probes used in the nCounter analysis.

**Table S2:** Calculated phases, amplitudes and periods of the Wt and *Pt*bHlH1a OE lines growing in 16L:8D. (P., Period; Ph., phase; Amp., amplitude; Std, standard deviation; t-test, t-test P-value between Wt and the indicated line).

| Data | N | Period | P.Std | P. t-test | Phase | Ph.Std | Ph. t-test | Amp. | Amp.Std | Amp. t-test |
| --- | --- | --- | --- | --- | --- | --- | --- | --- | --- | --- |
| Wt | 8 | 23.7 | 0.32 |  | 13.02 | 0.54 |  | 8.30E-01 | 1.20E-01 |  |
| OE-1 | 5 | 23.37 | 0.51 | 0.168894675 | 15.93 | 1.83 | 0.001213017 | 5.60E-01 | 2.60E-01 | 0.02447126 |
| OE-2 | 6 | 23.38 | 0.48 | 0.159824505 | 15.19 | 1.52 | 0.002642208 | 6.10E-01 | 7.00E-02 | 0.001606661 |
| OE-3 | 4 | 23.89 | 0.22 | 0.323356278 | 13.94 | 0.51 | 0.017556508 | 8.40E-01 | 1.20E-01 | 0.841165049 |

**Table S3:**
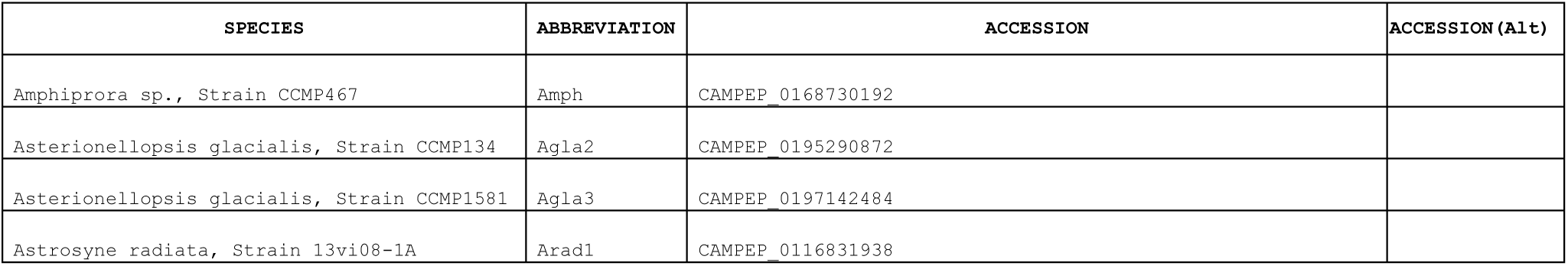
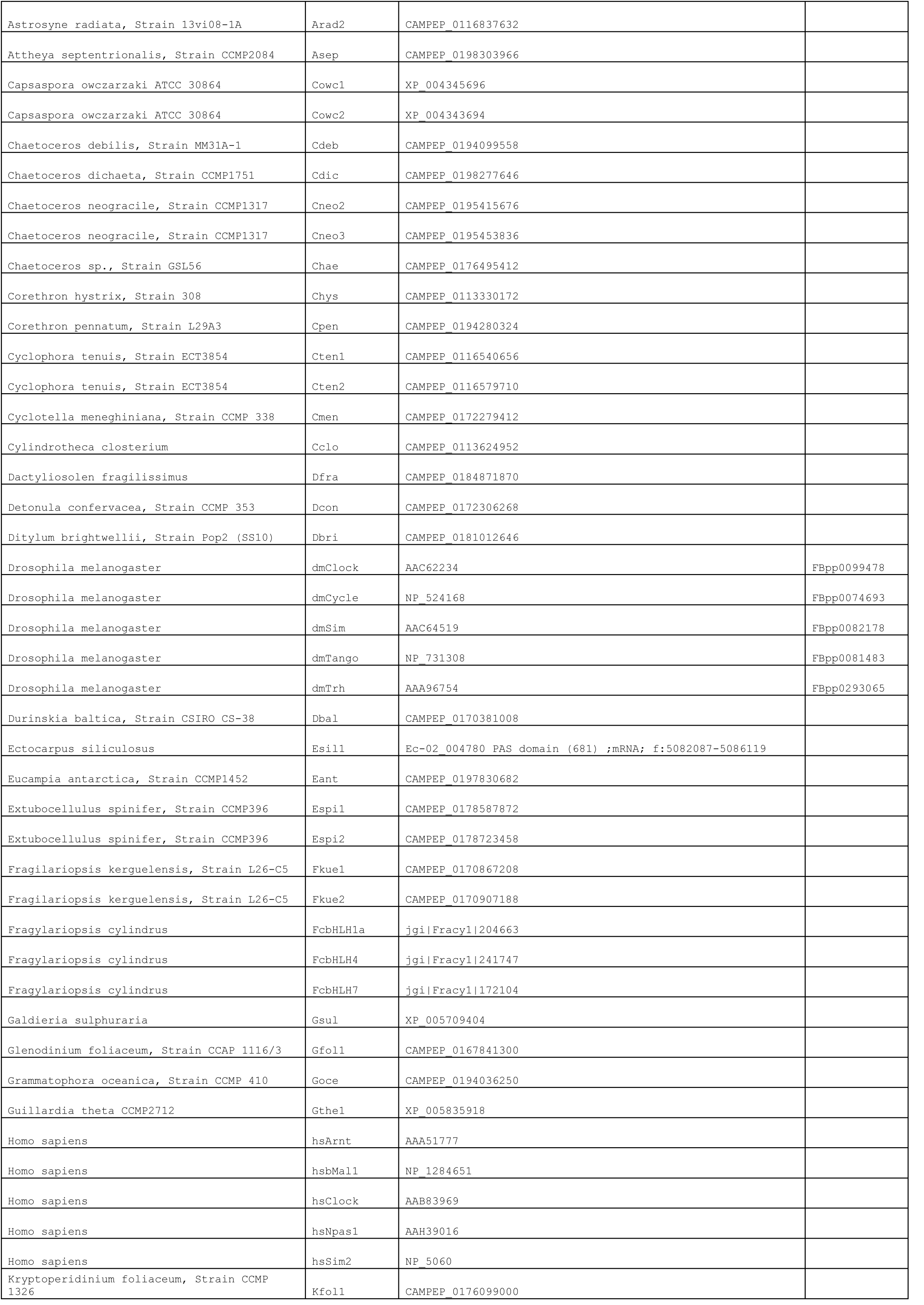
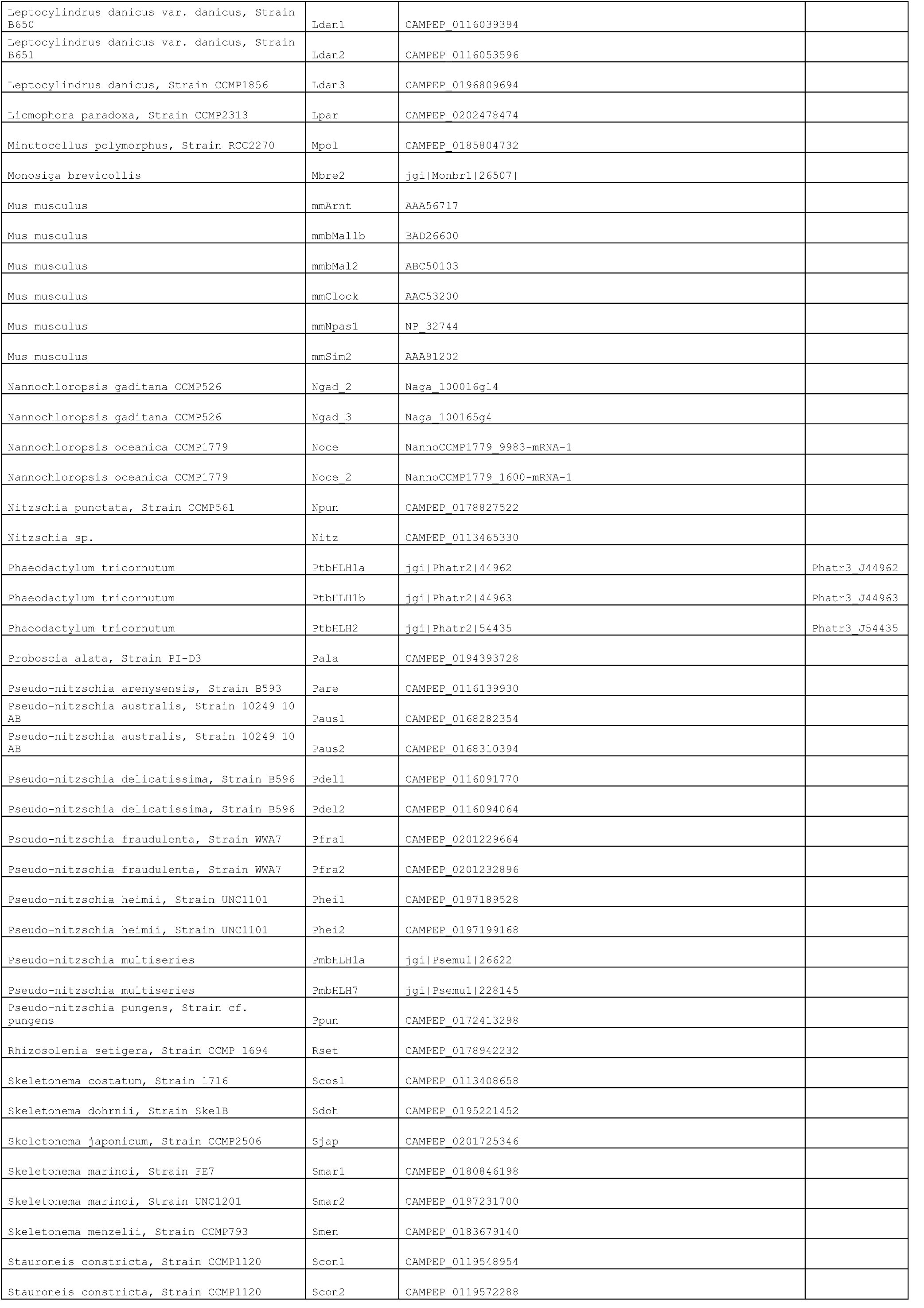
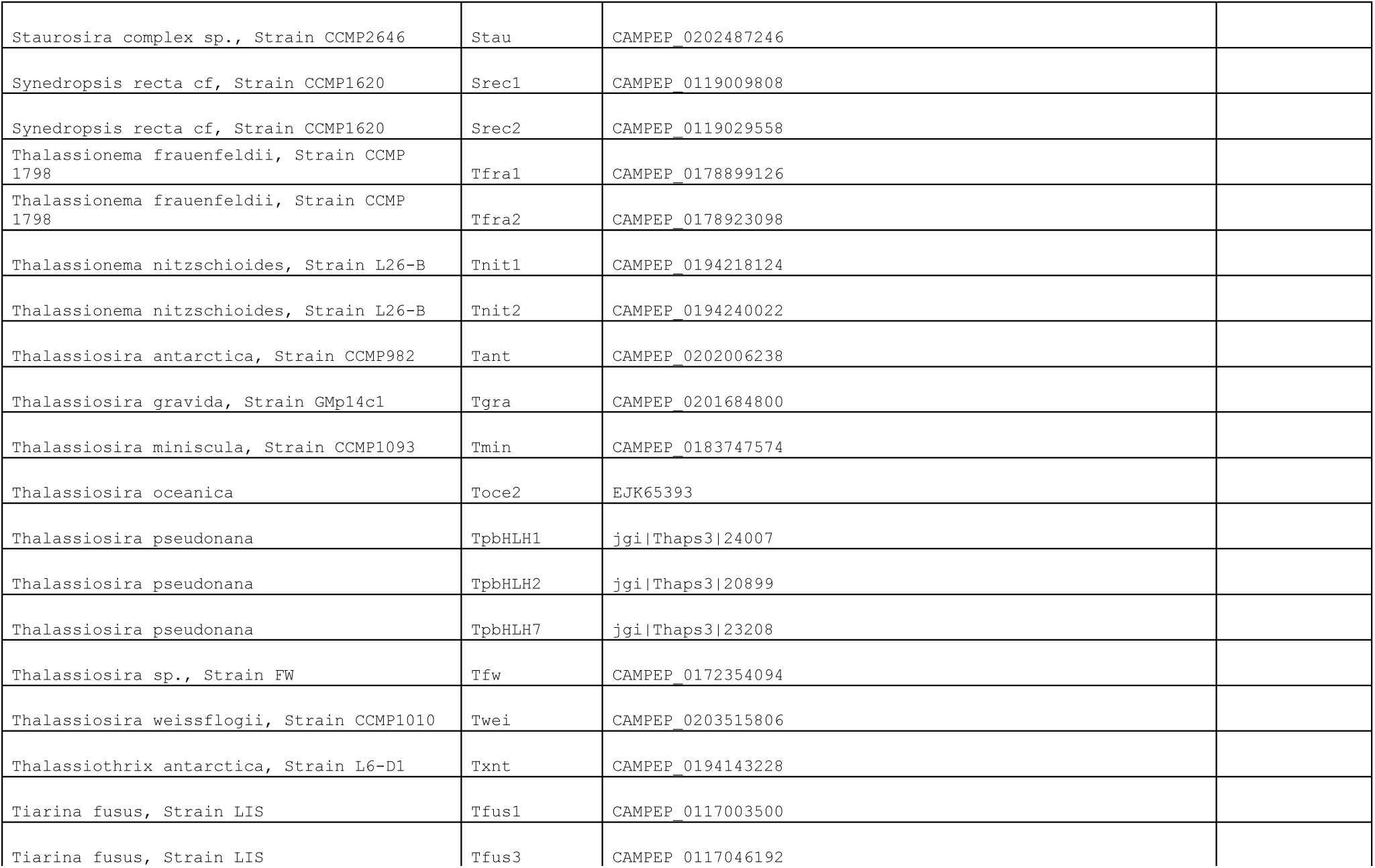
Accession numbers of the proteins utilized in the bHLH-PAS phylogenetic analysis.

**Table S4:**
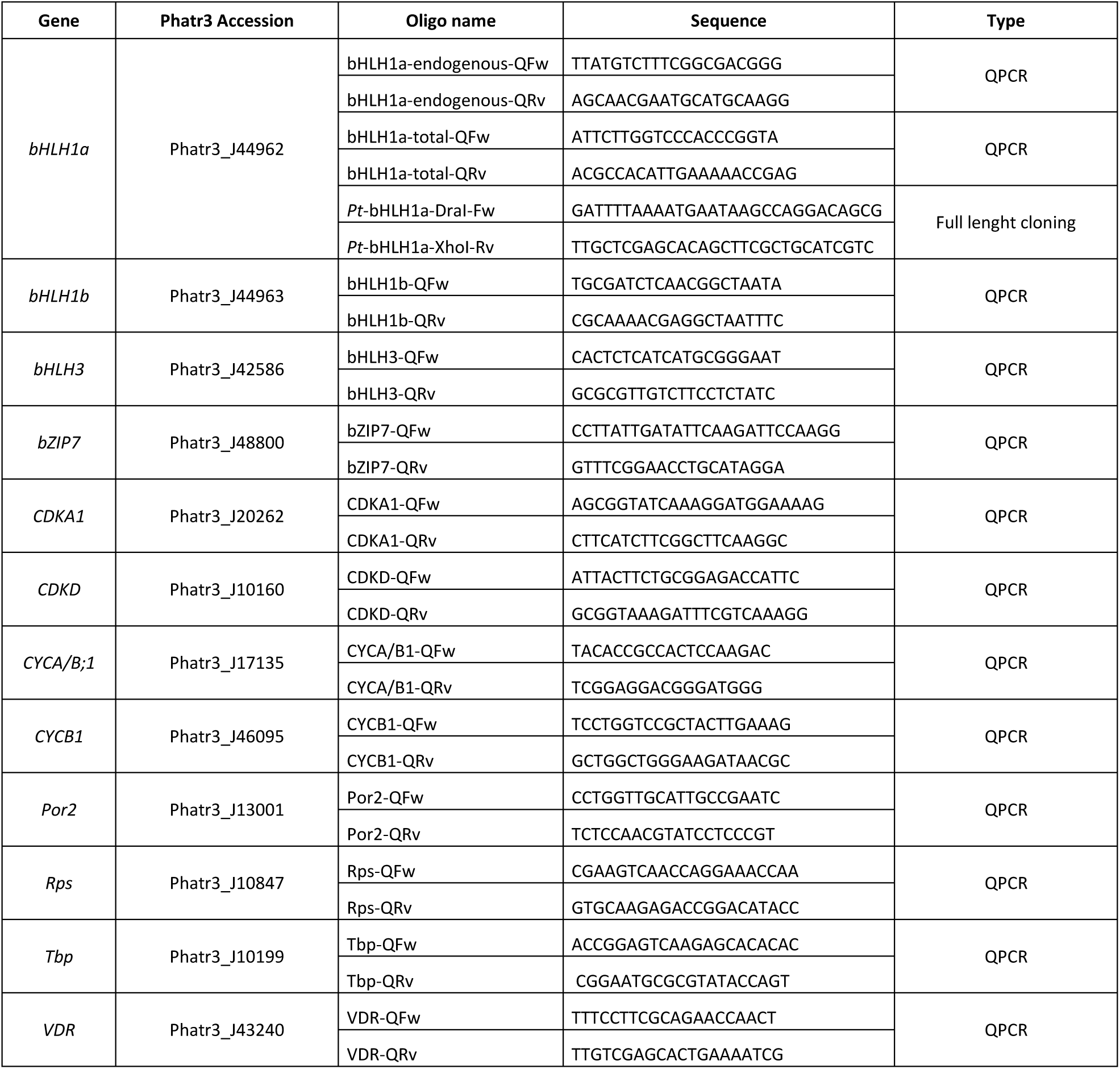
List of the oligonucleotides used in this work.

## REFERENCES

1. Pittendrigh CS (1993) Temporal Organization: Reflections of a Darwinian Clock-Watcher. Annual Review of Physiology 55(1):17–54.

2. Dodd AN, et al. (2005) Plant circadian clocks increase photosynthesis, growth, survival, and competitive advantage. Science 309(5734):630–633.

3. Woelfle MA, Ouyang Y, Phanvijhitsiri K, & Johnson CH (2004) The adaptive value of circadian clocks: an experimental assessment in cyanobacteria. Curr Biol 14(16):1481–1486.

4. Dunlap JC (1999) Molecular bases for circadian clocks. Cell 96(2):271–290.

5. Bell-Pedersen D, et al. (2005) Circadian rhythms from multiple oscillators: lessons from diverse organisms. Nat Rev Genet 6(7):544–556.

6. Chisholm SW (1981) Temporal Patterns of Cell-Division in Unicellular Algae. Can B Fish Aquat Sci (210):150–181.

7. Moulager M, et al. (2007) Light-dependent regulation of cell division in Ostreococcus: evidence for a major transcriptional input. Plant physiology 144(3):1360–1369.

8. Ragni M & D’Alcala MR (2007) Circadian variability in the photobiology of Phaeodactylum tricornutum: pigment content. J Plankton Res 29(2):141–156.

9. Naylor E (2010) Chronobiology of marine organisms (Cambridge University Press, Cambridge; New York) pp x, 242 p.

10. Tessmar-Raible K, Raible F, & Arboleda E (2011) Another place, another timer: Marine species and the rhythms of life. Bioessays 33(3):165–172.

11. Ottesen EA, et al. (2013) Pattern and synchrony of gene expression among sympatric marine microbial populations. P Natl Acad Sci USA 110(6):E488–E497.

12. Brierley AS (2014) Diel vertical migration. Current Biology 24(22):R1074–R1076.

13. Hafker NS, et al. (2017) Circadian Clock Involvement in Zooplankton Diel Vertical Migration. Current Biology 27(14):2194–2201.

14. Field CB, Behrenfeld MJ, Randerson JT, & Falkowski P (1998) Primary production of the biosphere: integrating terrestrial and oceanic components. Science 281(5374):237–240.

15. Malviya S, et al. (2016) Insights into global diatom distribution and diversity in the world’s ocean. Proc Natl Acad Sci U S A 113(11):E1516–1525.

16. Baldauf SL (2003) The Deep Roots of Eukaryotes. Science 300(5626):1703–1706.

17. Bowler C, et al. (2008) The Phaeodactylum genome reveals the evolutionary history of diatom genomes. Nature 456(7219):239–244.

18. Moustafa A, et al. (2009) Genomic footprints of a cryptic plastid endosymbiosis in diatoms. Science 324(5935):1724–1726.

19. Flori S, et al. (2017) Plastid thylakoid architecture optimizes photosynthesis in diatoms. Nat Commun 8:15885.

20. Raible F & Falciatore A (2014) It’s about time: Rhythms as a new dimension of molecular marine research. Mar Genomics 14:1–2.

21. Bailleul B, et al. (2010) An atypical member of the light-harvesting complex stress-related protein family modulates diatom responses to light. P Natl Acad Sci USA 107(42):18214–18219.

22. Allen AE, et al. (2011) Evolution and metabolic significance of the urea cycle in photosynthetic diatoms. Nature 473:203.

23. Goss R & Lepetit B (2015) Biodiversity of NPQ. Journal of Plant Physiology 172:13–32.

24. Morrissey J, et al. (2015) A novel protein, ubiquitous in marine phytoplankton, concentrates iron at the cell surface and facilitates uptake. Curr Biol 25(3):364–371.

25. McQuaid JB, et al. (2018) Carbonate-sensitive phytotransferrin controls high-affinity iron uptake in diatoms. Nature 555(7697):534–537.

26. Ashworth J, et al. (2013) Genome-wide diel growth state transitions in the diatom Thalassiosira pseudonana. P Natl Acad Sci USA 110(18):7518–7523.

27. Chauton MS, Winge P, Brembu T, Vadstein O, & Bones AM (2013) Gene regulation of carbon fixation, storage, and utilization in the diatom Phaeodactylum tricornutum acclimated to light/dark cycles. Plant physiology 161(2):1034–1048.

28. Smith SR, et al. (2016) Transcriptional Orchestration of the Global Cellular Response of a Model Pennate Diatom to Diel Light Cycling under Iron Limitation. PLoS Genet 12(12):e1006490.

29. Zones JM, Blaby IK, Merchant SS, & Umen JG (2015) High-Resolution Profiling of a Synchronized Diurnal Transcriptome from Chlamydomonas reinhardtii Reveals Continuous Cell and Metabolic Differentiation. Plant Cell 27(10):2743–2769.

30. Huysman MJ, et al. (2013) AUREOCHROME1a-mediated induction of the diatom-specific cyclin dsCYC2 controls the onset of cell division in diatoms (Phaeodactylum tricornutum). Plant Cell 25(1):215–228.

31. Huysman MJ, et al. (2010) Genome-wide analysis of the diatom cell cycle unveils a novel type of cyclins involved in environmental signaling. Genome Biol 11(2):R17.

32. Matthijs M, et al. (2017) The transcription factor bZIP14 regulates the TCA cycle in the diatom Phaeodactylum tricornutum. The EMBO journal 36(11):1559–1576.

33. Coesel S, et al. (2009) Diatom PtCPF1 is a new cryptochrome/photolyase family member with DNA repair and transcription regulation activity. EMBO Rep 10(6):655–661.

34. Kewley RJ, Whitelaw ML, & Chapman-Smith A (2004) The mammalian basic helix-loop-helix/PAS family of transcriptional regulators. Int J Biochem Cell Biol 36(2):189–204.

35. Rayko E, Maumus F, Maheswari U, Jabbari K, & Bowler C (2010) Transcription factor families inferred from genome sequences of photosynthetic stramenopiles. New Phytol 188(1):52–66.

36. Fortunato AE, et al. (2016) Diatom Phytochromes Reveal the Existence of Far-Red-Light-Based Sensing in the Ocean. Plant Cell 28(3):616–628.

37. Jaubert M, Bouly JP, Ribera D’Alcala M, & Falciatore A (2017) Light sensing and responses in marine microalgae. Curr Opin Plant Biol 37:70–77.

38. Hunsperger HM, Ford CJ, Miller JS, & Cattolico RA (2016) Differential Regulation of Duplicate Light-Dependent Protochlorophyllide Oxidoreductases in the Diatom Phaeodactylum tricornutum. PLoS One 11(7):e0158614.

39. Fortunato AE, Annunziata R, Jaubert M, Bouly JP, & Falciatore A (2015) Dealing with light: the widespread and multitasking cryptochrome/photolyase family in photosynthetic organisms. J Plant Physiol 172:42–54.

40. Oliveri P, et al. (2014) The Cryptochrome/Photolyase Family in aquatic organisms. Mar Genomics 14:23–37.

41. Banerjee A, et al. (2016) Allosteric communication between DNA-binding and light-responsive domains of diatom class I aureochromes. Nucleic Acids Res 44(12):5957–5970.

42. Young MW & Kay SA (2001) Time zones: a comparative genetics of circadian clocks. Nat Rev Genet 2(9):702–715.

43. Botebol H, et al. (2015) Central role for ferritin in the day/night regulation of iron homeostasis in marine phytoplankton. Proc Natl Acad Sci U S A 112(47):14652–14657.

44. Russo MT, Annunziata R, Sanges R, Ferrante MI, & Falciatore A (2015) The upstream regulatory sequence of the light harvesting complex Lhcf2 gene of the marine diatom Phaeodactylum tricornutum enhances transcription in an orientation-and distance-independent fashion. Mar Genomics 24 Pt 1:69–79.

45. Yan J, Ma Z, Xu X, & Guo AY (2014) Evolution, functional divergence and conserved exon-intron structure of bHLH/PAS gene family. Mol Genet Genomics 289(1):25–36.

46. Thiriet-Rupert S, et al. (2016) Transcription factors in microalgae: genome-wide prediction and comparative analysis. BMC Genomics 17:282.

47. Bhadra U, Thakkar N, Das P, & Bhadra MP (2017) Evolution of circadian rhythms: from bacteria to human. Sleep Med 35:49–61.

48. Merrow M, et al. (2006) Entrainment of the Neurospora circadian clock. Chronobiol Int 23(1-2):71–80.

49. Noordally ZB & Millar AJ (2015) Clocks in algae. Biochemistry 54(2):171–183.

50. Matsuo T, et al. (2003) Control mechanism of the circadian clock for timing of cell division in vivo. Science 302(5643):255–259.

51. Bunger MK, et al. (2000) Mop3 is an essential component of the master circadian pacemaker in mammals. Cell 103(7):1009–1017.

52. Gekakis N, et al. (1998) Role of the CLOCK protein in the mammalian circadian mechanism. Science 280(5369):1564–1569.

53. Taylor BL & Zhulin IB (1999) PAS domains: internal sensors of oxygen, redox potential, and light. Microbiol Mol Biol Rev 63(2):479–506.

54. Archibald JM (2015) Endosymbiosis and Eukaryotic Cell Evolution. Current biology 25(19):R911–921.

55. Guillard RRL (1975) Culture of Phytoplankton for Feeding Marine Invertebrates. In Culture of Marine Invertebrate Animals, D.R. Smith and M.H. Chanley, eds. Springer US.

## References

1. Zielinski T, Moore AM, Troup E, Halliday KJ, & Millar AJ (2014) Strengths and limitations of period estimation methods for circadian data. PLoS One 9(5):e96462.

2. Agier N & Fischer G (2016) A Versatile Procedure to Generate Genome-Wide Spatiotemporal Program of Replication in Yeast Species. Methods Mol Biol 1361:247–264.

3. Geiss GK, et al. (2008) Direct multiplexed measurement of gene expression with color-coded probe pairs. Nature biotechnology 26(3):317–325.

4. Chauton MS, Winge P, Brembu T, Vadstein O, & Bones AM (2013) Gene regulation of carbon fixation, storage, and utilization in the diatom Phaeodactylum tricornutum acclimated to light/dark cycles. Plant physiology 161(2):1034–1048.

5. Saeed AI, et al. (2003) TM4: a free, open-source system for microarray data management and analysis. Biotechniques 34(2):374–378.

6. Siaut M, et al. (2007) Molecular toolbox for studying diatom biology in Phaeodactylum tricornutum. Gene 406(1-2):23–35.

7. Keeling PJ, et al. (2014) The Marine Microbial Eukaryote Transcriptome Sequencing Project (MMETSP): Illuminating the Functional Diversity of Eukaryotic Life in the Oceans through Transcriptome Sequencing. Plos Biol 12(6).

8. Katoh K & Standley DM (2013) MAFFT multiple sequence alignment software version 7: improvements in performance and usability. Mol Biol Evol 30(4):772–780.

9. Kumar S, Stecher G, & Tamura K (2016) MEGA7: Molecular Evolutionary Genetics Analysis Version 7.0 for Bigger Datasets. Mol Biol Evol 33(7):1870–1874.

10. Darriba D, Taboada GL, Doallo R, & Posada D (2011) ProtTest 3: fast selection of best-fit models of protein evolution. Bioinformatics 27(8):1164–1165.

11. Miller MA, Pfeiffer, W., and Schwartz, T. (2010) Creating the CIPRES Science Gateway for Inference of Large PhylogeneticTrees. In Proceedings of the Gateway Computing Environments Workshop (GCE) (New Orleans, LA):1–8.

